# Decoding the olfactory map: targeted transcriptomics link olfactory sensory neurons to glomeruli

**DOI:** 10.1101/2021.09.13.460128

**Authors:** Kevin W. Zhu, Shawn D. Burton, Maira H. Nagai, Justin D. Silverman, Claire A. de March, Matt Wachowiak, Hiroaki Matsunami

**Affiliations:** Department of Molecular Genetics and Microbiology, Duke University Medical Center; Durham, North Carolina, 27710, USA; Department of Biological Sciences, Lehigh University; Bethlehem, Pennsylvania, 18015, USA; Department of Neurobiology, University of Utah School of Medicine; Salt Lake City, Utah, 84112, USA; College of Information Science and Technology, Pennsylvania State University; University Park, Pennsylvania, 16802, USA; Department of Statistics, Pennsylvania State University; University Park, Pennsylvania, 16802, USA; Department of Medicine, Pennsylvania State University; Hershey, Pennsylvania, 17033, USA; Institute for Computational and Data Science, Pennsylvania State University; University Park, 16802, USA; Department of Neurobiology, Duke University Medical Center; Durham, North Carolina, 27710, USA; Duke Institute for Brain Sciences, Duke University; Durham, North Carolina, 27710, USA

## Abstract

Sensory processing in vertebrate olfactory systems is organized across olfactory bulb glomeruli, wherein axons of peripheral sensory neurons expressing the same olfactory receptor co-terminate to transmit receptor-specific activity to central neurons. Understanding how receptors map to glomeruli is therefore critical to understanding olfaction. High-throughput spatial transcriptomics is a rapidly advancing field, but low-abundance olfactory receptor expression within glomeruli has previously precluded high-throughput mapping of receptors to glomeruli. Here we combined spatial sectioning along the anteroposterior, dorsoventral, and mediolateral axes with target capture enrichment sequencing to overcome low-abundance target expression. This strategy allowed us to spatially map 86% of olfactory receptors across the olfactory bulb and uncover a relationship between OR sequence and glomerular position.

**ONE SENTENCE SUMMARY:** Targeted enrichment of olfactory receptor mRNA in olfactory bulb sections determines spatial positions for murine glomeruli.

## MAIN TEXT

The organization of axonal projections from olfactory sensory neurons (OSNs) in the olfactory epithelium (OE) to glomeruli on the olfactory bulb (OB) forms the mammalian olfactory map (*1–4*). In the mouse, each canonical OSN expresses a single olfactory receptor (OR) or trace amine-associated receptor (TAAR) allele from a repertoire of over 1000 OR and TAAR genes. Insights into the organization of the olfactory map were first obtained using in situ hybridization, where OR transcript probes indicated the convergence of OSN axons into discrete structures called glomeruli on the OB surface, which range from 50 to 120 μm in diameter (*3, 5, 6*). Later, this organization was more clearly visualized through the use of gene-targeted mouse lines, which demonstrated that glomeruli are formed from the axonal convergence of OSNs expressing the same OR gene (*7*). Together these studies established the convergence of homotypic OSN axons to stereotyped glomeruli whose positional variability ranges from 75 to 270 μm depending on OR identity (*5, 8*). Because each glomerulus represents a single OR and a single odorant can bind multiple ORs, odor signals detected in the OE are transformed into a map of glomerular activity on the OB (*9–13*).

To date, glomerular positions for only ~3% of mouse ORs are available, and further progress has been stymied due to the low-throughput, laborious, and time-consuming aspects of currently available methodologies for mapping each OR in the expansive murine repertoire (*7, 8, 13–29*). Furthermore, the ability to compare locations between multiple glomeruli is limited among these studies due to the lack of reference landmarks on the OB and differences between methodologies. A more efficient approach for mapping OR axon projections to OB glomeruli would serve to generate a more comprehensive and informative map that would serve as a foundation for further functional studies of odor coding and processing. In this study, we demonstrate that target capture enrichment on spatial samples from the OB enables detection of low-abundance OR and TAAR mRNA in the axon termini of OSNs. Using this approach, we map 86% of the 1118 ORs and TAARs along the anteroposterior, mediolateral, and dorsoventral axes and combine these data to generate a three-dimensional model of glomerular positions with a precision of 141 μm. We examine the relationship between OR sequence and OB position, identify the set of ORs and TAARs expressed within dorsal glomeruli accessible to functional imaging, and generate gene-targeted mouse lines for two dorsal glomerular ORs amenable to functional characterization in vivo.

## RESULTS

### Targeted capture consistently enriches OR transcripts

Previous studies have detected low levels of OR mRNA in OSN axon terminals, identifying the glomerular positions for specific ORs within histological sections (*5, 23, 27*). To quantify OR and TAAR transcripts in the OB we first performed conventional RNA-Seq on whole-OB tissue from a mouse at postnatal day 21, an age when olfactory glomeruli are fully developed and finalized in their stereotyped positions (*30, 31*). Quantification of 25.7 million reads identified 410/1118 (36.7%) intact ORs at an average abundance of 0.06 transcripts per million (TPM) (median OR TPM = 0, 6/15 (40%) TAARs with TPM above 0, mean TAAR TPM = 0.077, median TAAR TPM = 0) (Fig. 1, B and C), confirming the low abundance of OR mRNA in OSN axon terminals.

**Fig. 1.**
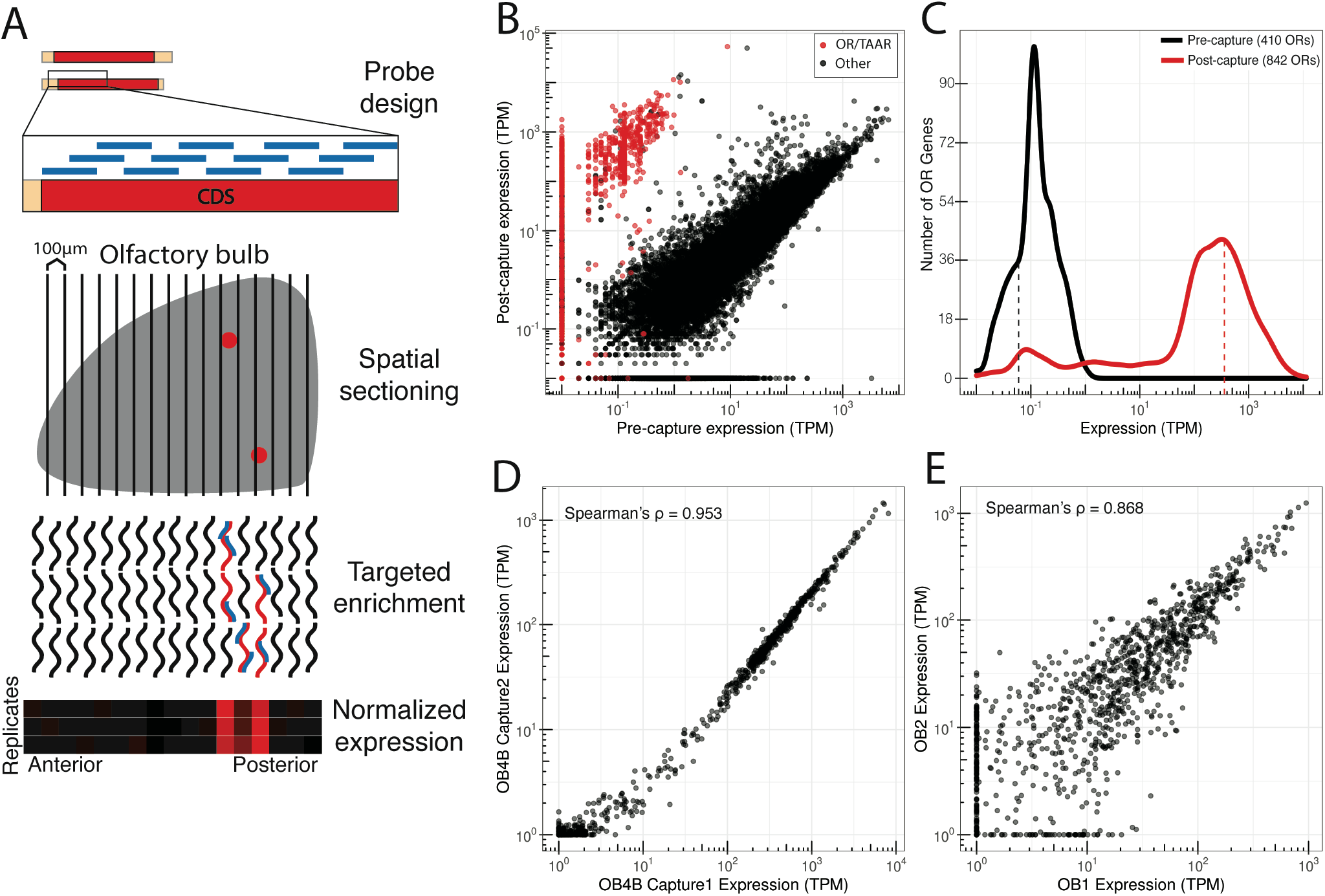
Target capture sequencing consistently enriches OR transcripts. (**A**) Methodological overview for targeted enrichment of OR and TAAR transcripts from OB sections. Briefly, RNA is extracted from OB tissue and used for cDNA synthesis and library preparation. Capture probes are designed against coding sequences (CDS) of interest, enabling enrichment of target genes following binding and washing steps. RNA-Seq of enriched libraries allows for high-throughput positional analyses when combined with systematic tissue sectioning. (**B**) Pre- and post-capture normalized abundances (transcripts per million; TPM) of intact OR and TAAR genes (red) and other intact genes (black) from a whole OB. (**C**) Frequency distribution of OR gene abundances pre- and post-capture from a whole OB. (**D**) Technical replicates demonstrating OR and TAAR gene abundances from independent capture enrichments of the same whole-OB RNA. (**E**) Biological replicates demonstrating OR and TAAR gene abundances from capture enrichment of whole-OB RNA samples from different animals.

To enrich sampling for OR transcripts, we designed a target capture array against chemosensory receptor gene families primarily targeting ORs and TAARs (*32*). We applied this target capture array to the previously sequenced OB library and identified 842 of the 1118 ORs (75.3%) and 10 of the 15 TAARs (66.7%) (Fig. 1, A to C). Following targeted capture, these ORs were present in a set of 27.7 million reads at an average abundance of 360.27 TPM (median = 106.74 TPM, mean TAAR TPM = 871.66, median TAAR TPM = 64.62) resulting in a mean fold enrichment of 6005X (mean TAAR fold enrichment = 11320X) (Fig. 1, B and C). Spearman’s rho for OR and TAAR transcript abundances between uncaptured and captured samples was 0.71 (*P* < 2.2 x 10^-16^). Further, four sets of independently captured technical replicates from two different OBs (two distinct spike-in mRNAs for RNA subsamples from each of the two bulbs, and two technical replicates per subsample) exhibited a Spearman’s rho of 0.95 (*P* < 2.2 x 10^-16^) (Fig. 1D and fig. S1D). The fold enrichment and correlation between pre- and post-capture samples indicates the targeted capture approach enriches the majority of ORs and TAARs in a highly consistent fashion as evidenced by the technical replicate correlation. The mean pairwise Spearman’s rho for three biological replicate OBs was 0.83 (*P* < 2.2 x 10^-16^), indicating the relative abundance of OR and TAAR transcripts is conserved between individual animals (Fig. 1E and fig. S1E).

In summary, targeted capture consistently enriched OR and TAAR transcripts to levels that facilitate positional analysis. This encouraged us to conduct targeted capture of ORs and TAARs from sections of OB to determine which ORs were expressed in each section. We sectioned from three directions, dorsoventral (DV), anteroposterior (AP), and mediolateral (ML). We note that these three axes are not precisely orthogonal and not perfectly concordant with the corresponding reference features such as DV zonal boundaries and medial surface of the OB.

### Expression of ORs and TAARs in dorsoventral OB sections correlates with OE expression zones

Pioneering investigation established that ORs are expressed in overlapping, continuous zones of the OE along the DV axis (*33, 34*). This zonal OE organization further correlates with DV glomerular positions of OR expression in the OB (*7, 23, 35*), with more recent studies leveraging multiplexed assays and transcriptomics to map an expanded number of ORs to more specific OE zones (*36, 37*). To comprehensively assess the relationship of OE-OB DV zonal organization of OR expression, we collected 100 μm sequential sections along the OB DV axis for targeted transcriptomics to determine which ORs were expressed in each section. Canonically, each OR is expressed in two glomeruli per bulb, both of which are expected to be located in similar positions along the DV axis. If enriched OR sequences are from OSN axon terminals, we expect that each OR would be abundant in spatially clustered sections which reflect the OE DV position from which the axons originate.

After weighting and normalization across individual mice, the localization pattern of each OR and TAAR was limited to a single spatial cluster in a series of neighboring sections for a majority of the capture-enriched transcripts (Fig. 2, A, B, and C). Uniform Manifold Approximation and Projection (UMAP) (*38*) visualization of data from the three DV replicate mice (22 sections per replicate) placed sequential sections from replicate animals in an ordered, non-clustered path, indicating that spatially related sections have similar transcriptional profiles (Fig. 2D). Replicate heatmaps were similar to each other (mean RVadj = 0.1803), which supports the stereotyped targeting of glomeruli to local domains (*39, 40*). To assess concordance of OB and OE positions along the DV axis, we constructed an expression-weighted mean DV position for each OR from the average of all three DV replicate mice, which we found correlated with the published OE DV positions of each OR (Spearman’s rho of mean position and OE index = 0.775, *P* < 2.2 x 10^-16^, Spearman’s rho = −0.690, *P* < 2.2 x 10^-16^) (Fig. 2E and fig. S2A) (*36, 37*). Along the DV axis, we found dorsal OE ORs, Class I ORs, and TAARs primarily located in the dorsal OB sections, while Class II ORs were distributed evenly along the DV axis, in agreement with previous mapping studies (dorsal vs ventral *P* = 4.292 x 10^-95^, Class I vs Class II *P* = 5.912 x 10^-20^, TAAR vs OR *P* = 0.0001, Mann-Whitney U-test) (Fig. 2F) (*29*).

**Fig. 2.**
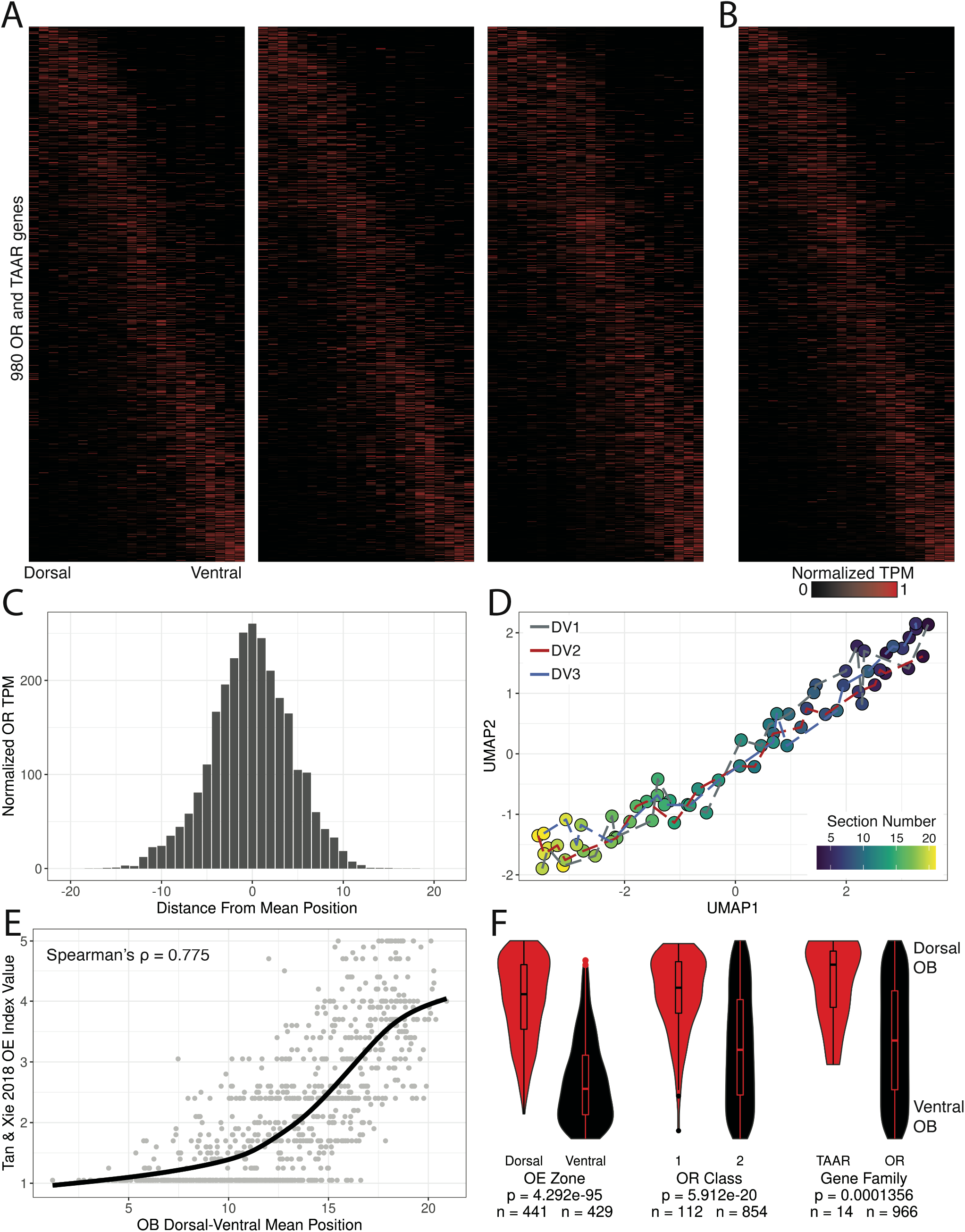
Glomerular OR expression correlates with OE expression along the DV axis. (**A**) Heatmaps for 980 ORs and TAARs across 22 DV sections sorted by mean position of expression from three replicate mice. (**B**) Merged representation of DV replicates. Ordering of genes (Y axis) is consistent across all heatmaps. (**C**) Distribution of normalized TPM (maximum observed value = 1, minimum observed value = 0) for all 980 ORs and TAARs from position of mean expression. (**D**) UMAP projection of 22 DV sections from all three replicates. (**E**) Line is loess smoothed regression of OE DV index from Tan and Xie, *Chem*. *Senses* 2018 across DV mean positions from our targeted spatial data. (**F**) Distribution of ranked DV mean positions for the 980 ORs and TAARs by OE zone, OR class, and gene family. Statistic is Mann-Whitney U-test.

### Anteroposterior spatial sections reflect stereotyped targeting of mirror-symmetric glomeruli

Using the same approach as for the DV axis, we examined 100-μm sections along the AP axis of the OB to further assess the precision and reproducibility of our method and the stereotypical patterning of OR glomeruli. Prior studies have shown that each OR typically has two glomeruli located in distinct, yet spatially linked AP and mediolateral (ML) positions, as each OB is organized into half bulbs along a non-orthogonal symmetry line (*8, 41, 42*). This symmetry typically leads to more posterior positioning of the medial glomerulus for an OR. However, in cases where the target location of an OR is close to the symmetry line, both glomeruli may appear in the same AP plane or only a single glomerulus may form (*43*). Based on these studies, we hypothesized that a majority of ORs would exhibit a bimodal expression pattern along the AP axis.

Spatial expression patterns of the normalized transcript abundances for 967 ORs and 14 TAARs were consistent across replicates, with stereotyped AP glomerular positions across OBs (*8, 44*). When sorted by position of mean expression, ORs primarily displayed two peaks of expression, consistent with published studies for labeled ORs displaying the medial glomerulus in a more posterior location relative to the lateral glomerulus (Fig. 3, A and B, and fig. S3A) (*8*). Compared to the distribution of normalized OR expression across the DV axis, ORs along the AP axis were distributed bimodally (Fig. 3C). Similar to the DV axis, UMAP projections of gene expression values from the six AP replicate mice (23 sections per replicate) revealed correlated expression patterns and AP positions across each OB (Fig. 3D). Across the AP axis, we found Class I ORs biased to the anterior set of sections, while TAARs tended to be localized to the central portion of the axis (Class I vs Class II *P* = 2.017 x 10^-26^, TAAR vs OR *P* = 0.7882, Mann-Whitney U-test) (Fig. 3E), consistent with previous studies examining glomeruli labeled in gene-targeted mice (*29, 45*). We also examined our data for concordance against the set of 32 ORs cloned from the anterior, middle, and posterior sections of an OB from Nakashima et al. (*46*). The ORs cloned from the anterior and middle OB (n = 13) had significantly more anterior mean positions than ORs cloned from the posterior OB (n = 18) (anterior + middle OB cloned ORs vs posterior OB cloned ORs *P* = 0.001, Mann-Whitney U-test) (Fig. 3F and fig. S4A). We further divided these cloned ORs across dorsal OE (n = 11) and ventral OE (n = 20) zones and found that both sets displayed concordance with our AP data, with ORs cloned from the anterior and middle OB positions having a lower AP mean position than ORs cloned from the posterior OB (dorsal OE: anterior and middle OB cloned ORs vs posterior OB cloned ORs *P* = 0.1636, ventral OE: anterior and middle OB cloned ORs vs posterior OB cloned ORs *P* = 0.0117, Mann-Whitney U-test) (fig. S4, B and C).

**Fig. 3.**
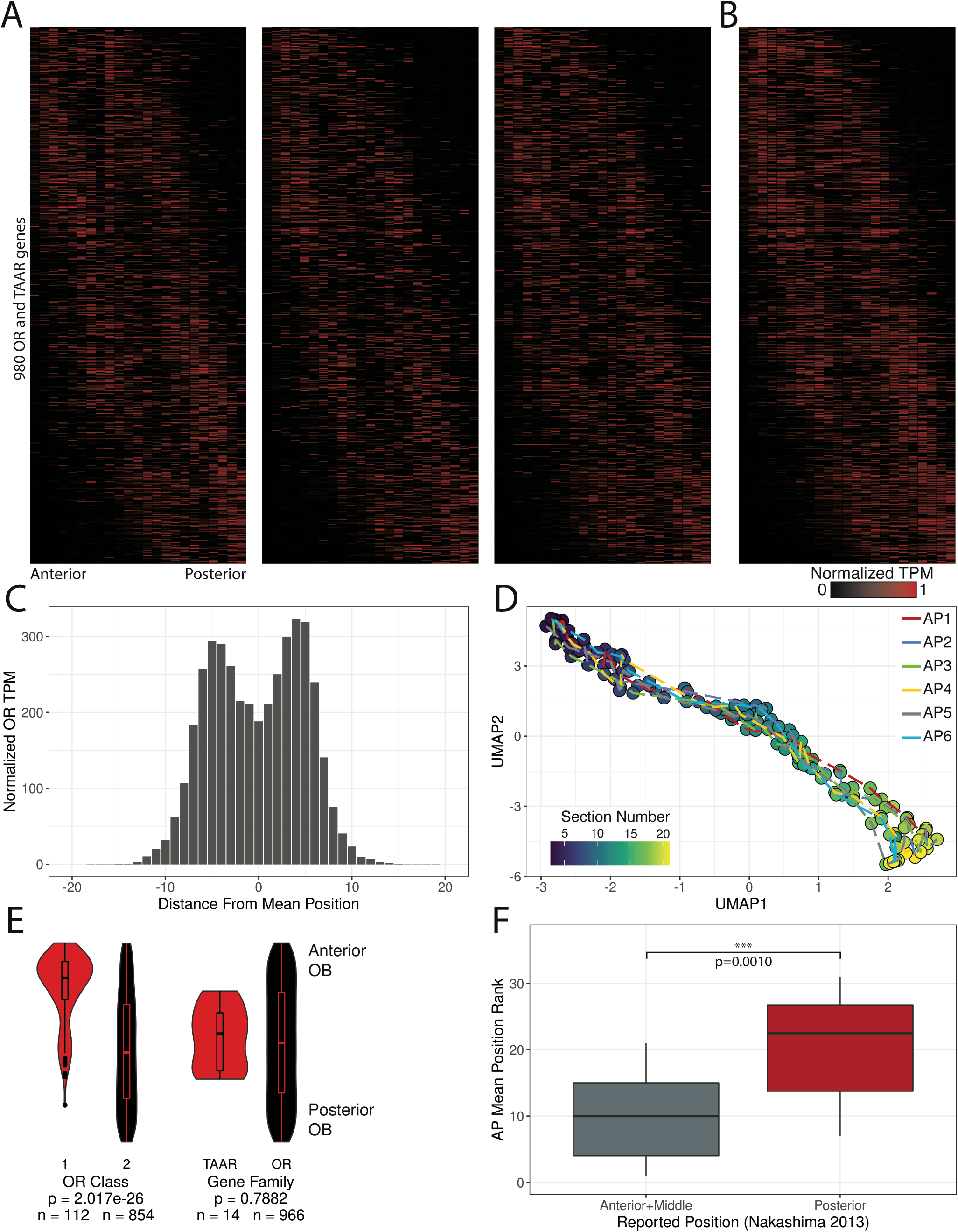
Glomerular OR expression is bimodally distributed along the anteroposterior axis. (**A**) Heatmaps for 980 ORs and TAARs across 23 AP sections sorted by mean position of expression from three replicate mice. (**B**) Merged representation from A. Ordering of genes is consistent across all heatmaps. (**C**) Distribution of normalized TPM (maximum observed value = 1, minimum observed value = 0) for all 980 ORs and TAARs from position of mean expression. (**D**) UMAP projection of 23 AP sections across all six replicates. (**E**) Distribution of ranked AP mean positions for the 980 ORs and TAARs by OR class and gene family. Statistic is Mann-Whitney U-test. (**F**) Distribution of the ranked AP mean position for the set of ORs cloned from the anterior and middle OB positions vs the posterior OB position from Nakashima et al. *Cell*. 2013. Statistic is Mann-Whitney U-test.

### Relationship of OR sequence and OB position

Gene-targeting approaches have identified a handful of examples in which OR sequence similarity correlates with glomerular position proximity (*24, 47*). Our dataset with a majority of ORs assigned to specific AP positions allowed us to systematically interrogate whether ORs with similar sequences exhibit similar glomerular positions by computing pairwise alignments for all ORs. To assess this relationship in all dimensions, we additionally generated a ML dataset (3 replicates, 22 sections per replicate) (fig. S5, A to E). Due to the combined presence of Class I ORs, Class II ORs, and TAARs on the dorsal surface of the OB, we separated our analysis into three groups: 1) Class I dorsal OB ORs, 2) Class II dorsal OB ORs, and 3) Class II ventral OB ORs. To assess OR similarity, we defined an OR gene alignment score threshold of 567, which corresponds approximately to the 40% identity threshold used to classify OR genes as belonging to a family (fig. S6A) (*48*). When comparing OR pairs above (≥567) and below (<567) the family-level threshold, we found both Class II dorsal and ventral ORs below the family level threshold displayed significant lower mean interglomerular distances across the AP, DV, and ML axes, suggesting topological relationship between the glomerular positions and family-level OR similarities for Class II ORs (Fig. 4A). In contrast, Class I ORs did not display a consistent relationship between sequence similarity and mean expression position along the DV and ML axes (median AP distance: 1.42 for ≥567 and 2.18 for <567 *P* < 2.2e-16, median DV distance: 2.59 for ≥567 and 2.44 for <567 *P* = 0.004, median ML distance: 2.95 for ≥567 and 2.67 for <567 *P* = 1.53e^-5^, Mann-Whitney U-test) (fig. S6B).

**Fig. 4.**
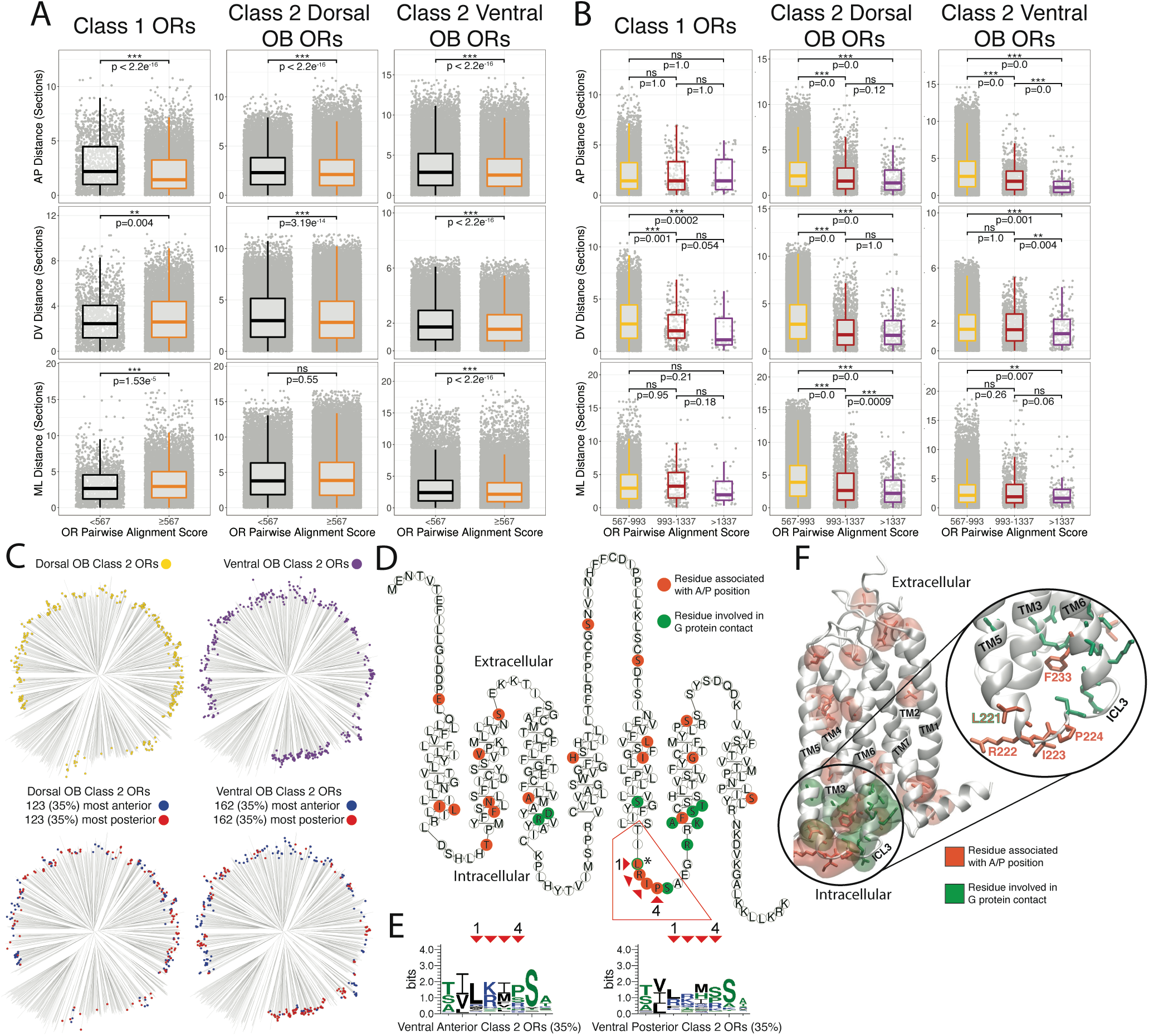
OR sequence similarity correlates with glomerular position more strongly among ventral than dorsal Class II ORs. (**A**) Pairwise comparisons between mean position distance and OR protein alignment score for all ORs separated by an alignment score cutoff relating to OR percent identity for family level classification with ORs below the family level score cutoff (<567) in black and above the family level score cutoff (≥567) in orange for the AP (top), DV (middle), and ML dimensions (bottom). Statistic is Mann-Whitney U-test. For Class I, n in “<567” = 1778, n in “≥567” = 8934. For Class II dorsal ORs, n in “<567” = 19346, n in “≥567” = 87256. For Class II ventral ORs, n in “<567” = 26718, n in “≥567” = 37544. (**B**) Pairwise comparisons between mean position distance and OR protein alignment score for ORs above family level classification (40%, 567) separated by an alignment score cutoffs relating to OR percent identity for subfamily level classification (60%, 993) and highly similar ORs (80%, 1337) for the AP (top), DV (middle), and ML dimensions (bottom). ORs falling within the range of family to subfamily level ORs (567-993) are displayed in yellow, subfamily to highly similar level ORs (993-1337) are displayed in red, and above the highly similar level OR score cutoff (>1337) in magenta. Statistic is Mann-Whitney U-test. For Class I, n in “567-993” = 8632, n in “993-1337” = 248, n in “>1337” = 54. For Class II dorsal ORs, n in “567-993” = 85144, n in “993-1337” = 1830, n in “>1337” = 282. For Class II ventral ORs, n in “567-993” = 36204, n in “993-1337” = 1126, n in “>1337” = 214. (**C**) Phylogenetic tree of all Class II dorsal OB ORs (top left, n = 354), all Class II ventral OB ORs (top right, n = 464), the most anterior (35%, blue) and most posterior (35%, red) Class II ORs from the dorsal (bottom left) and ventral OB (bottom right) DV zones. (**D**) Snakeplot of the Class II OR consensus protein sequence; orange residues have significantly different physicochemical properties for ventral, anterior or posterior, Class II ORs compared to all ventral Class II ORs. Green residues indicate residues known to be involved in Class A GPCR activation through contact with the G protein (* indicates the single residue which was identified as being both associated with G protein contact and identified as significantly different for ventral, anterior, Class II ORs). (**E**) Protein sequence logos for the four identified intracellular loop 3 residues associated with ventral Class II anterior/posterior ORs depicting the conservation of specific amino acid residues at each position. Red arrows indicate the specific residue within the Class II OR consensus snakeplot (C) and the corresponding position in the sequence logo. (**F**) Homology model of the mouse Class II consensus OR. Transmembrane helices (TM) are numbered with residues associated with A/P positions (orange) and residues in contact with the G protein (green) depicted in licorice with transparent regions indicating the residue surface. Right, highlight of intracellular loop 3 (ICL3) where residues associated with A/P positions are intermingled with the residues in contact with the G protein.

We further examined this relationship by comparing ORs above the 40% (family level ORs), 60% (subfamily level ORs) and 80% (highly similar, used to define OR orthologs) protein identity thresholds (*48, 49*) using the pairwise alignment score thresholds 993 and 1337 respectively. Comparisons between family (567-993), subfamily (993-1337), and highly similar ORs (>1337) revealed similar results among the Class II ORs (Fig. 4B). Along the AP axis, both dorsal and ventral Class II ORs displayed progressively more similar glomerular positions in groups with higher sequence similarity with statistical significance except for one comparison (Fig. 4B and Table S5). Results for both dorsal and ventral Class II ORs along the DV and ML axes and for Class I ORs along all axes indicated that among the groups of related ORs, glomerular positions typically become more similar or do not change with increasing sequence similarity. Altogether our data generally agrees with a model in which overall similarities of ORs influence the relative glomerular locations.

We next sought to determine if any specific OR amino acid residues correlated with AP position. Due to the different relationships between sequence similarity and OB position for dorsal and ventral ORs, we examined Class II ORs using different cutoffs for the sets of the most anterior and most posterior ORs included in the analysis (20%, 27.5%, 35%) to identify amino acid residues correlating with AP position (Fig. 4C and fig. S7A, B, C). We identified 22 residues whose physicochemical properties differed from all ventral Class II ORs (Fig. 4D and fig. S8C and S9) (*50*). Notably, four consecutive residues that correlated with AP position were in the third intracellular loop, which has been shown to interact directly with the G protein during Class A GPCR activation (Fig. 4, D to F, and fig S8, A and B) (*51*). Additionally, the phenylalanine within the KAFSTCxSH motif is sandwiched between four residues involved in G-protein binding, (*52–54*). These findings indicate residues that are at or near the sites of G-protein interactions are critical in determining glomerular position, which is consistent with the hypothesis that ligand-independent basal activity of ORs influences glomerular targeting (*43, 46, 55–57*).

### A three-dimensional model of OR glomerular positions reflects established features

To systematically estimate OR glomerular positions across the entire OB, we merged our AP, DV, and ML datasets into a unified 3D model. UMAP plots of the mean position of ORs in each dimensional replicate indicated that ORs fall along a 3D axis oriented from anterior-dorsal-medial to posterior-ventral-lateral (fig. S10, A and B). To account for the location of OR glomeruli on the outer surface of the OB, we extracted coordinates from a diffusion tensor imaging (DTI) model of the mouse OB to represent the approximate glomerular layer that would have been sampled by each section along each dimension of our targeted transcriptomics data (*58*). We applied a Bayesian model that computes probability distributions for each OR in each voxel based on the isometric-log ratio transform of the normalized read counts from the corresponding set of intersecting AP, DV, and ML single-dimension samples (*59, 60*). These normalized counts were weighted by the proportion of TPM of total olfactory marker protein (OMP; expressed in all mature OSNs) originating from that position, normalized by the proportion of voxels located in that section. Finally, we developed an algorithm for the systematic reduction of positional probabilities across the whole OB surface into predicted OR glomerular positions which were then filtered to remove predictions with posterior median values below 0.0005 to account for ORs with low expression. Posterior median summaries for the resulting set of 709 ORs and TAARs in all voxels are freely viewable as interactive 3D maps at kanazian.shinyapps.io/obmap/.

We assessed our algorithm by comparing predictions for Class I and II ORs, dorsal and ventral OE ORs, and the set of ORs examined via transgenic mouse lines by Zapiec et al. (*8*). Predicted glomerular positions for Class I ORs, Class II ORs, and dorsal and ventral ORs were consistent with expectations based on OE zone, OR class, and our single-dimension data (dorsal vs ventral *P* = 8.26 x 10^-99^, Class I vs Class II *P* = 1.962 x 10^-12^, functional imaging surface enriched vs not-enriched *P* = 1.38 x 10^-18^ (see below), Mann-Whitney U-test) (Fig. 5, A and E and fig. S11, C, and D). Additionally, the distribution of predictions for the sets of Class I ORs and TAARs matched previously published findings for target domains (Fig. 5B and fig. 11). Our current model predicts glomerular positions for 700 ORs and 9 TAARs, with predicted positions outperforming randomly selected ORs from the same DV zone and medial/lateral side for the subset of ORs with known positions with a median error of 141 μm (Fig. 5, C and D) (*8*). The relative positions of the Olfr1377 and Olfr881 glomeruli in the gene-targeted mouse lines (see below) were also consistent with those predicted from the spatial transcriptomic data (Fig. 5C and figs. S13 and S14 fig. S12D), providing further support for our three-dimensional glomerular predictions based on the spatial transcriptomic data. In summary, we found our three-dimensional reconstruction of OR glomerular positions to be both in agreement with established OR spatial features and to show greater concordance with these established features than the sets of single-dimension target capture sequencing data alone. Collectively, our results thus provide the first large-scale, unified, and systematic database of OR glomerular positions for the mouse OB.

**Fig. 5.**
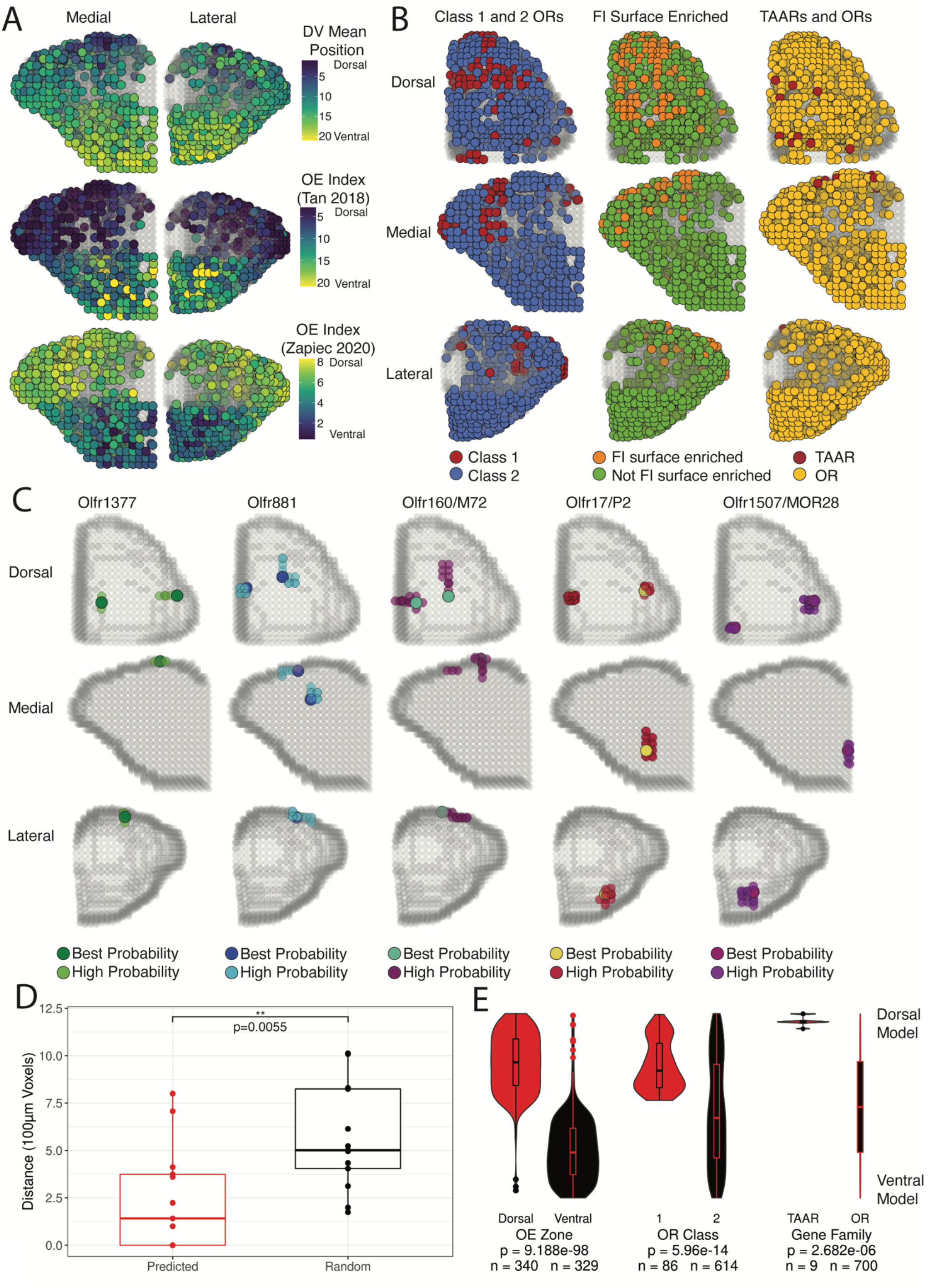
A three-dimensional model for OR glomerular positions from combined single-dimension targeted sequencing data. (**A**) Three-dimensional predictions for the 709 ORs and TAARs with DV mean positions (top) and with observed values in OE DV indices (middle and bottom). (**B**) Three-dimensional predictions for the 709 ORs and TAARs with colors revealing contrasting distributions of Class I vs Class II ORs, functional imaging surface enriched vs not enriched ORs, and TAARs vs ORs. (**C**) Three-dimensional predictions for labeled ORs generated in this study and from Zapiec and Mombaerts *PNAS* 2015. High probability positions indicate the set of adjacent voxels containing highly ranked probabilities for that OR with the best probability color indicates the voxel with the highest ranked probability within that cluster of voxels. (**D**) Distance between predicted OR position and OR positions determined from gene-targeted mouse lines compared to 50 random ORs from the same DV zone (15 ORs used for unusual zone) and side of the OB. Statistic is Mann-Whitney U-test. (**E**) Distribution of ranked mean DV model position for the best probability voxel in each predicted glomerulus for all 709 ORs and TAARs for OE zone, OR Class, and gene family. Statistic is Mann-Whitney U-test.

### Identification of ORs within the dorsal functional imaging window

We sought to validate and extend our findings by examining specific ORs that map to glomeruli on the dorsal-central OB surface, which has been extensively characterized by functional imaging in vivo (*9, 11, 12, 44, 61*). To define the set of ORs accessible under standard functional imaging approaches, we collected tissue samples from OBs from C57BL6 mice (2 male, 2 female). Each OB was dissected into two parts, one containing the approximate dorsal-central imaging area and the other containing the remainder of the OB (fig. S12A). These 16 samples were processed for target capture sequencing and differential expression analysis to identify ORs enriched in the functional imaging area.

A total of 121 ORs, including 27 Class I ORs and 94 Class II ORs, were consistently enriched in the imaging surface (FDR ≤ 0.05 and LogFC > 0) (Fig. 5B, Fig. S12, B, E and F), with 96% of these ORs known to localize in dorsal OE zones (*36*). We also found nine of the 15 TAARs enriched in the imaging surface, with no TAARs enriched in the remaining OB tissue (fig. S12G), consistent with previous functional imaging of some TAAR glomeruli (*45*). To anatomically and functionally validate our expression analysis, we chose two ORs as targets for the generation of receptor-tagged gene-targeting mouse lines, based on their enrichment in the functional imaging area (Olfr881: FDR = 0.007, LogFC = 2.55 and Olfr1377: FDR = 0.021, LogFC = 2.14) (fig. S12B) and their robust response to the odorant 4-methylacetophenone in pS6-IP RNA-Seq experiments (Olfr1377: FDR = 3.43e^-26^, LogFC = 3.18, Olfr881: FDR = 7.37e^-23^, LogFC = 3.18). (fig. S12C) (*62*). Using *Easi*-CRISPR (*63*), we inserted *IRES-mKate2* cassettes following the CDS of each OR to create Olfr1377-IRES-mKate2 and Olfr881-IRES-mKate2 mice, in which OSNs expressing either Olfr1377 or Olfr881 also express the cytosolic fluorescent marker mKate2, labeling cell bodies in the OE and glomeruli in the OB (fig. S12D). Whole-mount confocal imaging of the OB in gene-targeted mice revealed mKate2-labeled glomeruli within the dorsal-central OB surface (Fig. 6A and fig. S13, A and B).

**Fig. 6.**
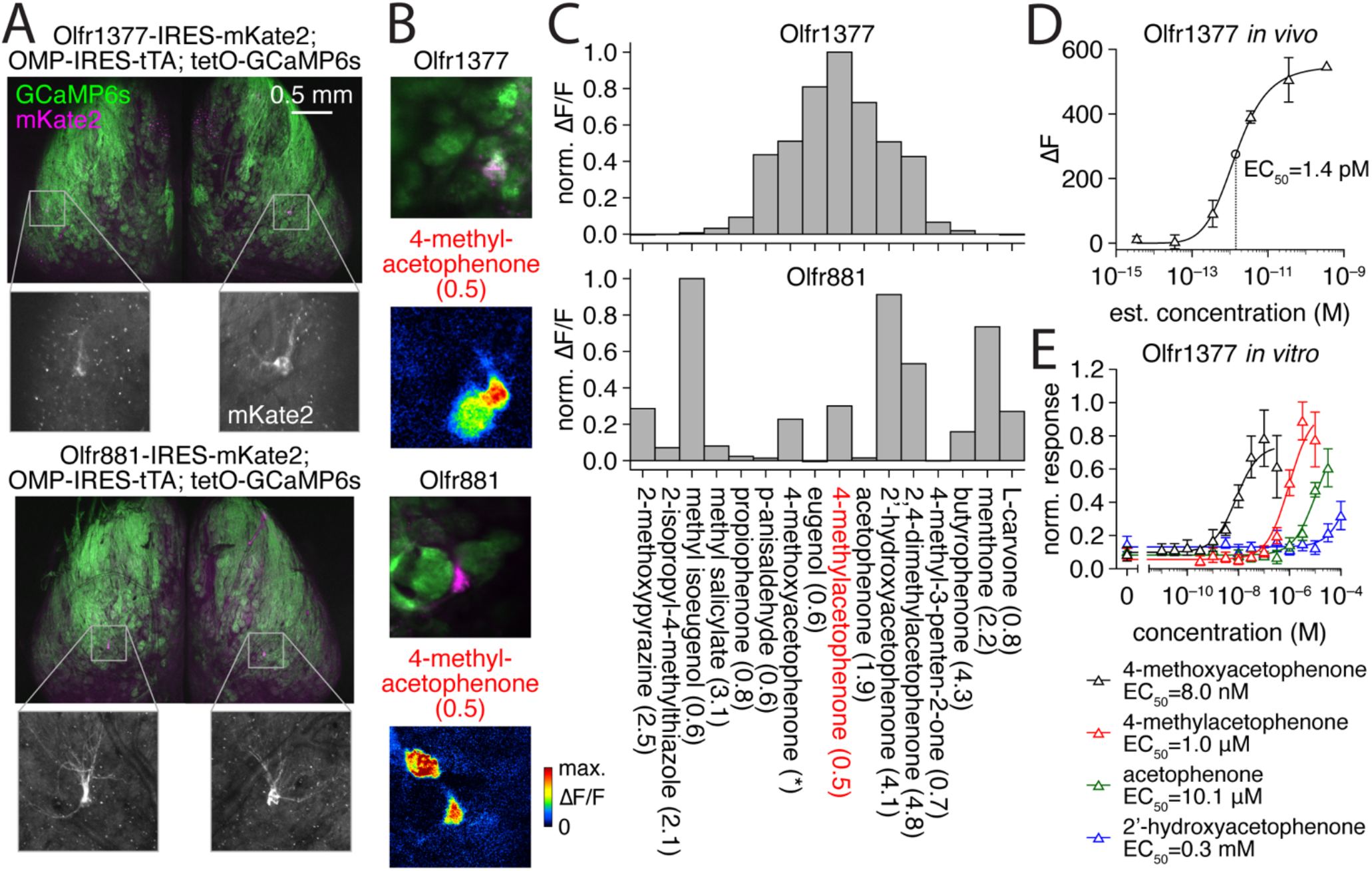
Deorphanization of Olfr1377 and Olfr881. (**A**) Whole-mount confocal maximal intensity projection of compound heterozygous Olfr1377-IRES-mKate2; OMP-IRES-tTA; tetO-GCaMP6s and Olfr881-IRES-mKate2; OMP-IRES-tTA; tetO-GCaMP6s mice following two-photon functional imaging. (**B**) Baseline fluorescence and GCaMP6s ΔF/F responses to 2-s presentation of 4-methylacetophenone during two-photon functional imaging of the heterozygous lines in (A). Estimated concentration of delivered odorant (in nM) provided in parentheses here and elsewhere. Olfr1377 response map scaled from 0-80% ΔF/F. Olfr881 response map scaled from 0-30% ΔF/F. (**C**) Spectra of Olfr1377 and Olfr881 glomerular ΔF/F responses (sorted by Olfr1377 response magnitude) to a subset of ligands detected within a larger odorant panel. Odorants presented at an estimated concentration on the order of 10^0^ nM, with the exception of 4-methoxyacetophenone (~3.5 nM for Olfr881; ~3.5 x 10^-3^ nM for Olfr1377). (**D**) In vivo concentration-response function of the Olfr1377 glomerulus to 4-methoxyacetophenone. (**E**) In vitro concentration-response function of the Olfr1377 receptor to an array of cyclic ketone odorants.

Examination of additional whole-mount epifluorescence images allowed us to further assess position and variance of both Olfr1377 and Olfr881 (fig. S14, Fig. 5C). Consistent with the singular expression of most ORs as two mirror-symmetric glomeruli, both gene-targeted mouse lines labeled two glomeruli per OB. The lateral Olfr1377 glomerulus (n = 10) was positioned centrally along on the AP axis and central-laterally within the ML axis imaging area, while the medial glomerulus (n = 6) was more posterior, ventral, and medial. The lateral Olfr881 glomerulus (n = 9) was positioned centrally along the ML axis and relatively posterior within the imaging area, while the medial glomerulus (n = 8) displayed a more variable position across the medioposterior quarter of the dorsal surface. The lateral Olfr1377 glomerulus displayed nearly twice (196.8%) the positional variance than the lateral Olfr881 glomerulus while the medial Olfr1377 glomerulus was distributed in an area nearly half the size (52.8%) of its Olfr881 counterpart.

### Functional imaging of dorsal ORs

Expression of long-wavelength mKate2 as an OR-specific marker allowed for functional characterization of Olfr1377 and Olfr881 glomeruli by crossing the generated mouse lines to OSN-specific driver lines expressing a GCaMP Ca^2+^reporter. For maximal imaging sensitivity, we crossed each mKate2 line to the OMP-IRES-tTA driver line (*64*) and the tetO-GCaMP6s reporter line (*65*). In the resulting triple crosses, we readily located the lateral mKate2-tagged glomeruli on the dorsal functional imaging surface (Fig. 6A) and imaged odorant-evoked GCaMP6s signals from these and neighboring glomeruli using dual-wavelength two-photon imaging in anesthetized mice. Consistent with our pS6-IP in vivo data (fig. S12C), both Olfr1377 and Olfr881 exhibited robust responses to low concentrations of 4-methylacetophenone, with Olfr1377 exhibiting a stronger response than Olfr881 (Fig. 6, B and C). In addition to 4-methylacetophenone, we also tested a large odorant panel including multiple cyclic ketones structurally related to 4-methylacetophenone, as well as more diverse odorants, all at relatively low concentrations. From this panel, we identified multiple new, high-affinity ligands for each OR, including many cyclic ketones, and ultimately uncovered overlapping but distinct response spectra for Olfr1377 and Olfr881 (Fig. 6C, fig. S15C). For example, Olfr1377 showed strong responses to p-anisaldehyde, acetophenone, and the aliphatic ketone 4-methyl-3-penten-2-one, while Olfr881 proved unresponsive to these odorants.

Interestingly, Olfr1377 (but not Olfr881) exhibited an exceptionally strong response to 4-methoxyacetophenone, with a brief (2 s) presentation of ~0.4 nM 4-methoxyacetophenone eliciting long-term activation and desensitization (fig. S15, A and B). Additional concentration screening suggested an in vivo response threshold of ~10^-13^ M 4-methoxyacetophenone for the Olfr1377 glomerulus (Fig. 6D and fig. S15D). To complement our in vivo imaging and pS6-IP RNA-Seq analyses and further evaluate the sensitivity of Olfr1377 to 4-methylacetophenone and 4-methoxyacetphenone, we additionally expressed Olfr1377 in Hana3a cells and examined luciferase responses to different concentrations of 4-methoxyacetophenone, 4-methylacetophenone, 2’-hydroxyacetophenone, and acetophenone (*66, 67*). Reinforcing our in vivo results, Olfr1377 responded to all four odorants (Fig. 6E). Moreover, Olfr1377 responded to 4-methoxyacetophenone at just 1 nM and saturated at 100 nM in vitro, a response orders-of-magnitude more sensitive than the response to any other odorant tested. Collectively, these findings thus identify 4-methoxyacetophenone-Olfr1377 as a ligand-receptor interaction with exceptionally high affinity.

## DISCUSSION

Olfactory receptor neurons in *Drosophila* predominantly express a single OR out of a repertoire of 62 OR genes and project their axons to one of ~50 distinct glomeruli on each antennal lobe (*68, 69*). These glomeruli have been individually linked to specific ORs, allowing for a complete picture of how sensory information is organized within the first olfactory relay (*69–71*). This key map has served as a foundation for critical studies regarding sensory processing, olfactory receptor neuron-targeting factors, the propagation of information to higher-order neurons, and the cellular composition of the antennal lobe (*72–75*). In contrast, the mammalian system involves over an order of magnitude more receptors and lacks such a comprehensive and specific mapping.

With over 1000 ORs projecting to approximately 1800 glomeruli arrayed over the three-dimensional surface of the mouse OB, the experimental challenge of mapping ORs to glomeruli is profound (*76*). To date, such mapping in the mouse has relied primarily on the creation of gene-targeted animals or in situ hybridization of radiolabeled probes. Further, the lack of finescale landmarks within the OB and differences between methods used to identify glomerular positions has led to relatively limited comparable positional information, with single studies examining just six ORs by in situ hybridization and five by fluorescent markers (*8, 26*). Given the low-throughput of these previous assays and the large size of the mouse OR repertoire, the ability of the field to comprehensively assess the genetic and logical organization of the olfactory map has been limited.

Here, we demonstrate a unique application of target capture sequencing, enriching low-abundance target transcripts found in the axon terminals of sensory neurons and comparing post-enrichment abundances between samples. We applied this method to 100 μm sections along the three cardinal axes to circumvent a longstanding problem in olfaction: resolving the receptor organization of the olfactory map. Our high-throughput approach generated the first repertoire-scale dataset for OR glomerular positions and enabled the creation of a three-dimensional positional estimate for the glomeruli of a majority of the OR repertoire with an approximate precision within ranges previously observed for positional variability between animals. The scale of our study enabled us to verify the longstanding observation that OE zones map directly to OB glomeruli along the dorsoventral axis. Additionally, our dataset allowed us to begin interrogating how OR sequence is related to OB target position, an important aspect for the formation of the olfactory map. We found that, in general, OR sequence similarity correlates with OB target position for both classes of ORs and for both the dorsal and ventral hemispheres of the OB. Targeting along the AP axis has further been hypothesized to be associated with ligand-independent basal activity of the OR protein located at the axon terminal membrane. We defined a set of amino acid residues correlated with the most-anterior and most-posterior ventral Class II ORs. Intriguingly, many of these residues were proximal to domains involved in G-protein coupling, which raises the possibility for involvement in modulation of basal activity levels, which would in turn influence targeting along the AP axis. Mutational studies testing a diverse set of ORs in both in vitro and in vivo paradigms will likely be needed to determine the exact effect of each residue as basal activity is dependent on the OR, secondary structure interactions, and the membrane the receptor is embedded within.

While our data lacks single glomerulus-level spatial resolution, our three-dimensional model provides a probabilistic model of OR glomerular positions that leverages biological replicates and glomerular position stereotypy. We provide a visualization of the most likely voxel for a given glomerulus and high probability voxels indicating the potential variance in position. We provide predictions for all 980 ORs and TAARs analyzed in our single-dimension heatmaps in an online application where we provide an interactive model with values and filters for probabilities of our predictions.

In developing this three-dimensional model, we also identified the set of ORs and TAARs present within the dorsal surface that is typically viewed in functional imaging studies, which, almost exclusively, lack information regarding the OR identity of the observed glomerular responses. Identification of this set, representing ~10% of the OR repertoire, will directly facilitate the in vivo deorphanization of ORs by guiding combined gene-targeting and functional imaging approaches – a strategy crucial for validating and fine-tuning complementary in vitro deorphanization screens (*13, 77*). Using this approach, we identified numerous ligands for two previously uncharacterized ORs - Olfr881 and Olfr1377 - including odorants activating both receptors (e.g., 4-methylacetophenone; methyl isoeugenol) as well as odorants selectively activating only Olfr881 (e.g., menthone) or Olfr1377 (e.g., p-anisaldehyde). In addition, we uncovered an exceptionally sensitive and long-lasting response of Olfr1377 to 4-methoxyacetophenone, with affinity paralleling the ultrasensitive detection of amines by TAARs (*77*). Of great interest, this latter result tentatively suggests that high-affinity odorant-receptor interactions are not exclusively limited to TAARs and physiologically important amines, but may exist for a broader collection of ORs and diverse odorants.

The projection of OSN axons from the OE to OB glomeruli requires exquisite accuracy and precision. The degree of inter-individual variation requires replicate animals to be sampled and limits the precision to which any approach can achieve, and our approach provides a level of precision that is comparable to the resolution limits set by stereotyped targeting. The mapping of OR glomerular positions to specific subdomains of the OB broadly serves to generate unique insights into how this early sensory relay center is organized and reflects our chemical environment. Joining ligand-receptor deorphanization data with our newly generated OR-OB positional data enables large-scale investigations into how representations of odor space relate to topographical features in the OB and facilitates the development of unified, systematic models for how chemical inputs are processed, interpreted, and transformed into odor-driven behaviors.

## ACKNOWLEDGMENTS

We thank members of the Matsunami lab for thoughtful discussions and feedback. Luis R. Saraiva (SIDRA Institute), Antonio Scialdone (Helmholtz Zentrum München), and Mayra L. Ruiz Tejada Segura (Helmholtz Zentrum München) provided helpful advice on mapping OR expression and spatial transcriptomics. We thank Sayan Mukherjee (Duke University) for guidance on the combination of single dimension transcriptomic data into a three dimensional model. We thank C. Ron Yu (Stowers Institute, Univ. of Kansas) for providing OMP-IRES-tTA mice. We thank Shelby Priest (Duke University) for assistance with critical reading and feedback.

## AUTHOR CONTRIBUTIONS

Conceptualization: KWZ, MW, HM

Methodology: KWZ, SDB, JDS, MW, HM

Investigation: KWZ, SDB, MHN, MW, HM

Visualization: KWZ, SDB, MHN, CAdM, MW, HM

Funding acquisition: KWZ, SDB, MW, HM

Project administration: MW, HM

Supervision: MW, HM

Writing – original draft: KWZ, SDB, MW, HM

Writing – review & editing: KWZ, SDB, MHN, JDS, CAdM, MW, HM

## DATA AND MATERIALS AVAILABILITY

All data are available in the main text or the supplemental materials.

