## Supplemental materials for "Decoding the olfactory map: targeted transcriptomics link olfactory sensory neurons to glomeruli"

### <u>SUPPLEMENTARY MATERIALS:</u>

### MATERIALS AND METHODS

#### *Code availability and statistics*

Relevant code has been made publicly available at https://github.com/kanazian/obmap. All statistical tests were performed in R. All Mann-Whitney U-tests were two-tailed and performed with continuity corrections using the wilcox.test function from the stats package. Dunn's Krukal Wallis test performed with Benjamini Hochberg false discovery rate correction using the dunn.test function from the dunn.test package. All box and whisker plots display the median value as the middle line, the first and third quartile as the hinges, and the farthest values within 1.5 inter-quartile range of the first or third quartile as the middle line as the whiskers. * indicates p value $\leq 0.05$, ** indicates p value $\leq 0.01$, *** indicates p value $\leq 0.001$.

#### *Animals*

Wild-type C56BL6 mice were purchased from the Jackson Laboratory and used for breeding. Olfr1377-IRES-mKate2 and Olfr881-IRES-mKate2 mice were generated by inserting *IRES-mKate2* cassettes downstream of the CDS of each OR using *Easi*-CRISPR (*63*). tetO-GCaMP6s mice (*65*) were purchased from the Jackson Laboratory (Stock No. 024742). OMP-IRES-tTA mice (*64*) were provided by C. Ron Yu (Stowers Institute, Univ. of Kansas). Mice were group housed with food and water available ad libitum and kept on a 12-h-light/dark cycle. All procedures for animal handling and tissue collection were approved by the Institutional Animal Care and Use Committees of Duke University and the University of Utah.

#### ***Sample acquisition and sequencing library preparation***

Pups were sacrificed between postnatal day 20 and 22. Whole brains with olfactory bulbs intact were dissected and placed in a solution of 3% Low Melting Point Agarose (American Bioanalytical) within an embedding mold (Polysciences Inc, Peel-A-Way R30) before placing on ice. Once solidified, the mold was removed and the agarose-embedded brain was prepared for vibratome sectioning by cutting away agarose to leave a triangular shape with the tip forming at the bulb on the surface that will be cut first for that specific direction. The triangular block was glued (Loctite 404) onto the vibratome stand (Leica VT1000S with Feather carbon blades), submersed in cold 1X HBSS (Gibco) and 100 µm sections serially cut. Sections were placed into 1.5mL tubes and stored at -80C. 400 µL of Buffer RLT (Qiagen) with 10% BME was added to each frozen tissue section before homogenization at 30,000RPM for 4 seconds using a mounted Biogen PRO200 with 5mm flat tip generator probe. The homogenizer probe was rinsed three times with DI water between samples. RNA was extracted using a Qiagen RNeasy kit according to the manufacturer’s instructions. cDNA synthesis was performed using a SMART-Seq v4 (Takara) kit according to the manufacturer’s instructions with 10 ng total RNA used for input in half-sized reaction volumes. 10 ng cDNA was used in the KAPA Hyperplus library construction kit following the manufacturer’s instructions. All quantifications were performed using Qubit.

#### ***Target capture sequencing and alignment***

We selected a set of ORs, TAARs, V1Rs, and V2Rs for inclusion in our assay based on the mouse genome annotation available at the time (GRCm38.p4 release M10). This set of transcripts was submitted to Roche Nimblegen for inclusion in a targeted enrichment probe panel. All sections from a single OB were uniquely indexed and combined in equal amounts to create a 1000 ng pool which was processed for target capture according to the manufacturer's protocol using a 20-hour hybridization period. Target capture library pools were sequenced on an Illumina NextSeq 500 Sequencing System in the 75SR or the HiSeq2500 in 50SR mode at the Duke Center for Genome and Biology shared core facility. Snakemake v3.5.5 (*78*) was used to process read files through alignment and quantification. STAR v2.7.0d (*79*) was used to generate a genome index using the primary assembly and comprehensive gene annotation file of the mouse genome (Gencode GRCm38.p6 release M25). Reads were aligned to this genome index using STAR with default options except for –quantMode TranscriptomeSAM which maps genome alignments to transcript coordinates. STAR output transcriptome SAM files were quantified using RSEM v1.3.1 (*80*) using default options to generate gene level counts for differential expression.

***Sample normalization***

TPM values for each sample from each replicate OB were weighted by the proportion of TPM of OMP for that section divided by the total TPM from all samples from the replicate. Weighted TPM was then normalized between 0 and 1 using the minimum and maximum values for each OR. Merged heatmaps were generated using the median normalized values at each position of the expression matrix for DV and ML heatmaps (n

= 3) and the median of four replicates plus the mean of value of the four replicates at that position for the AP heatmaps. Matrix of median values were then renormalized for each OR across samples. The following ORs were excluded from all analyses due to abnormally high expression values that potentially indicate ectopic expression of the OR within OB cell types: Olfr287, Olfr32, Olfr361, Olfr1033.

***Sequence and position analysis and identification of anterior and posterior OR associated residues***

Mouse OR protein sequences were downloaded from HORDE (*81*) and OR pairwise alignments were computed using the pairwiseAlignment function from the R package Biostrings using the BLOSUM62 substitution matrix, a gapOpening penalty of 11 and a gapExtension penalty of 1. Approximately corresponding pairwise alignment scores and percent identity values were extracted from a curve determined by calculating the mean percent identity value for three different pairs of ORs at each pairwise alignment score divisible by 100 using the Expasy SIM alignment tool (https://web.expasy.org/sim/) with parameters matching those used for pairwise alignment (data not shown). 1090 mouse OR protein sequences were aligned using Clustalx with manual adjustments as previously published (*62*). To identify residues that were more conserved by the 139 (30% set) most anterior, ventral, Class II ORs than by chance, we simulated distributions of Grantham distances for random subsets of 70 (50% of size of set) ventral Class II ORs for all pairwise combinations for each residue in the alignment. Random sampling was performed 1000 times. The per-residue mean Grantham distances were computed for all combinations of the 139 most anterior, ventral, Class II

ORs and used to find p-values under the null distribution after FDR correction. Alignment positions with gaps in more than 10% of OR sequences were excluded from the analysis. Analysis was repeated for all sets (20%, 27.5%, 35%) for both anterior and posterior Class II ORs. Visualization of the structural sequence for Olfr539 were generated using GPCRsnakeplotter (https://github.com/Yue-Jiang/snakeplotter). Sequence logos were created using WebLogo (http://weblogo.threeplusone.com/) using default settings.

***Differential expression analysis***

The R package, edgeR (v3.34.0), was applied to the sample count table for all genes in the reference (*82*). Common, trended, and tagwise dispersions were estimated separately and in sequence, respectively. For identification of ORs in the functional imaging surface and ORs responding to 4-methylacetophenone, results were subset to olfactory receptor genes (pseudogenes included) and new FDR values were calculated from the original p values determined using all genes.

***Homology model building***

The protocol follows a previously published method (*53*). Aligned protein sequences of 1092 mouse ORs were manually aligned to pre-aligned protein sequences of 11 GPCRS including bovine rhodopsin (PDB ID 1U19), human chemokine receptors CXCR4 (PDB ID 3ODU) and CXCR1 (PDB ID 2LNL), and human adenosine a2A receptor (PDB ID 2YDV) using Jalview (*83*). Four experimental GPCR structures (1U19, 3ODU, 2YDV, and 2LNL) were used as templates to build the Class II mouse consensus OR by homology modeling using Modeller. Five models were obtained and the one fulfilling several constraints (sufficiently large binding cavity, no large folded structure in extracellular loops, all TMs folded as α-helices, and a small α-helix structure between TM3 and TM4) was kept for further visualization.

***Three-dimensional model of OR glomeruli positions***

A DTI model of the mouse brain (*58*) was converted from .nii to .stl filetype and loaded in Blender. An array of cubes with the number of cubes per dimension matching the number of sections collected from each dimension was stretched in each dimension to encompass the entire OB in order to account for difference in age of mice. OB and cube structures were exported as separate .ply files and imported into R where a custom script was used to determine the position of cubes which contain OB surface polygons. Additional OB surface-containing cubes were included to remove gaps, model the medial surface, and account for a 100 μm wide glomerular layer within the outer surface of the OB. The final set of OB surface-containing cubes was exported as three-dimensional coordinates for use as a scaffold for the shape of the OB glomerular layer. The composition of glomeruli in each voxel along the OB surface was calculated as the weighted average of the composition obtained from sequencing each OB section that intersected that voxel weighted by the total number of surface voxels in each section. To account for sequencing noise, the composition of glomeruli in each sequence was estimated using a Bayesian multinomial-Dirichlet model with a Dirichlet prior parameter of 0.65 reflecting the weak prior knowledge that each section likely contained only a subset of all possible glomeruli in the OB. For each sequenced section, 1000 posterior samples from this model were obtained and used in subsequent calculations of the glomerular composition in each surface voxel. Custom functions were applied to determine positional clusters of high probability voxels for each gene based on the Bayesian posterior probability and the expected DV location of the OR based on class and Tan and Xie *Chem. Senses.* 2018 index identity. Briefly, each gene was designated as dorsal, ventral, unusual zone, or non-OR based on gene name, OR class, OE position, and OB mean position. Based on this DV designation, the highest probability voxel cluster (minimum 3 adjacent voxels) from the corresponding DV hemisphere on each side of the line of symmetry was picked. The probability of the medial and lateral clusters was then compared and used to define the region of mirror symmetry for the cluster with the lower probability with the lower probability cluster being repicked from a mirror symmetry field across the line of symmetry. If the medial cluster had a higher probability, the lateral cluster was repicked from a field that was from the lateral half and ranging from 6 sections anterior to 1 section posterior of the medial cluster. If the lateral cluster had a higher probability, the medial cluster was repicked from a field that was from the medial half and ranging from 6 sections posterior to 1 section anterior of the lateral cluster. Clusters that were found within 6 sections of the front or back of the model were repicked from fields between the end of the model and 6 sections inward. ORs classified as being in the unusual zone in the OE were predicted as a single large cluster within the anterior, ventral, central face of the OB. Non-OR genes were predicted using the same method as dorsal ORs with only the top probability voxel cluster being returned.

***pS6-IP RNA-Seq***

Three P21 mice of either sex were used per odorant or control condition. Animals were individually habituated in sealed containers for 1 h, then transferred to a new container containing a piece of filter paper enclosed in a cassette (Tissue-Tek) spotted with 10 μl of 4-methylacetophenone diluted at 1% (v/v) in distilled water or with 10 μl of distilled water (no odor control). After 1 hr of exposure, the animals were euthanized, then had their olfactory epithelium dissected in cold 25 ml of dissection buffer (1 × HBSS +$Ca^{2+}$, +$Mg^{2+}$ [Gibco], 2.5 mM HEPES pH 7.4, 35 mM glucose, 100 μg/ml cycloheximide, 5 mM sodium fluoride, 1 mM sodium orthovanadate, 1 mM sodium pyrophosphate, 1 mM beta-glycerophosphate). The dissected tissues were transferred to 1.35 ml of cold homogenization buffer (150 mM KCl, 5 mM $MgCl_2$, 10 mM HEPES pH 7.4, 100 nM Calyculin A, 2 mM DTT, 100 U/ml RNasin [Promega], 100 μg/ml cycloheximide, 5 mM sodium fluoride, 1 mM sodium orthovanadate, 1 mM sodium pyrophosphate, 1 mM beta-glycerophosphate, 1x protease inhibitor [Roche]). Bones were removed, and the tissue was mechanically dispersed three times at 250 rpm and nine times at 750 rpm using a homogenizer (Glas-Col). The homogenates were centrifuged at 4600 rpm for 10 min at 4°C in a 1.5 ml Lobind tube (Eppendorf), then the supernatants were collected into a new 1.5 ml Lobind tube and combined with a mixture of 90 μl 10% NP-40 and 90 μl 300 mM DHPC (Avanti Polar Lipids). The samples were centrifuged at 13000 rpm for 10 min at 4°C, then the supernatants were collected into a new 1.5 ml Lobind tube and incubated with 20 μl pS6 antibody (Cell Signaling #5364) for 1.5 hr at 4°C under constant rotation. During the last 30 minutes of incubation, 100 μl Protein A Dynabeads (Invitrogen) were prepared by washing three times with 900 μl of wash buffer 1 (150 mM

KCl, 5mM $MgCl_2$, 10 mM HEPES pH 7.4, 0.05% BSA, 1% NP-40). The samples were added to the beads and incubated under constant rotation for 1 hr at 4°C, followed by four washes with 700µl of wash buffer 2 (350 mM KCl, 5 mM $MgCl_2$, 10 mM HEPES pH 7.4, 1% NP-40, 2 mM DTT, 100 U/ml recombinant RNasin [Promega], 100 µg/ml cycloheximide, 5 mM sodium fluoride, 1 mM sodium orthovanadate, 1 mM sodium pyrophosphate, 1 mM beta-glycerophosphate). The samples were moved to room temperature at the final wash, and the elution was performed with 350 µl Buffer RLT (Qiagen). The RNA present in the eluate was purified with the RNeasy Micro kit (Qiagen).

***Heterologous cell expression***

The Dual-Glo luciferase assay was performed as described previously (*84*). Briefly, Hana3A cells (*85*) were plated on 96-well plates in MEM supplemented with 10% FBS. After 18-24 hr incubation at 37 °C and 5% $CO_{2,}$ cells were transfected with 100 ng/µL plasmids coding for Olfr1377 (5 µL/plate), muscarinic acetylcholine receptor M3R (2.5 µL/plate), RTP1S (5 µL/plate), pRL-SV40 (5 µL/plate) and CRE-luciferase (10 µL/plate) using Lipofectamine 2000 (Invitrogen). pCI vector (5 µL/plate) was used instead of Olfr1377 plasmid for the transfection of control cells. 18-24 h after the transfection, the culture medium was replaced with serial dilutions of the odorants in CD293 medium (GIBCO), then the plates were incubated for 3.5 hours at 37°C and 5% $CO_2$ without the lid. Luminescence was measured using a Polarstar Optima plate reader (BMG). Luminescence values (LV) were determined by subtracting basal luminescence (of an empty 96-well plate), then dividing firefly luminescence by renilla luminescence (to control for transfection efficiency and cell viability) as in the formula: LV = (firefly luminescence - 400)/(*Renilla* - 400). The LV values were normalized and plotted using GraphPad Prism 9 by establishing the minimum LV = 0 and the maximum LV = 1 and using the analysis log(agonist) vs. response (three parameters). CAS numbers for tested odorants are as follows: acetophenone - 98-86-2, 2-hydroxyacetophenone - 118-93-4, 4-methylacetophenone - 122-00-9, 4-methoxyacetophenone - 100-06-1.

***Functional imaging***

Adult mice (2-3 months old) of both sexes were anesthetized and prepared for functional imaging with 3 Hz artificial inhalation via a double-tracheotomy procedure as previously described (Burton Chem Senses 2019). Imaging was conducted using a resonant-scanning two-photon microscope (Sutter Instruments) coupled to a pulsed Ti:Sapphire laser (Mai Tai HP, Spectra Physics) tuned to 920 nm and a Fidelity-2 1070 nm laser (Coherent) and equipped with a 16X 0.8 N.A. water-immersion objective (Nikon), with emission separated by green (520/65 nm) and red (641/75 nm) filters and collected by GaAsP photomultipliers (Hamamatsu H10770B). Glomeruli were imaged at 15.5 Hz through thinned-bone windows. Odorants were presented using a custom olfactometer (Burton Chem Senses 2019) in pseudorandom order (except for 4-methoxyacetophenone concentration series), with 2 s delivery duration and variable inter-delivery intervals. Odorants were diluted in caprylic/capric medium chain triglycerides (C3465, Spectrum Chemical Mfg. Corp.).

For response maps and spectra (Fig. 6B and C), ΔF/F responses were calculated as the difference in mean fluorescence within 4 s following odorant onset from the 4 s preceding odorant onset (the baseline fluorescence), divided by the baseline fluorescence. For display, ΔF/F response maps were bilinearly interpolated by a factor of 2 and low-pass filtered (Gaussian, standard deviation = 1 pixel) to improve map resolution and reduce map noise, respectively. For the concentration-response function of the Olfr1377 glomerulus to 4-methoxyacetophenone (Fig. 6D), ΔF responses were calculated as the difference in mean fluorescence within 2.5 s following odorant onset from the baseline fluorescence. For the lowest three odorant concentrations (in which responses fully decayed before subsequent odorant presentations), baseline fluorescence was calculated as the mean fluorescence within the 1 s preceding odorant onset. For the highest three odorant concentrations (in which responses failed to fully decay before subsequent odorant presentation), ΔF responses were calculated using the baseline fluorescence observed during the final $3.5 \times 10^{-13}$ M odorant presentation. All ΔF/F and ΔF responses are averaged across three odorant presentations. All analysis was performed using custom code written in MATLAB (MathWorks).

***Whole-mount visualization***

Adult mice of both sexes were transcardially-perfused with PBS followed by 4% paraformaldehyde and postfixed overnight before brains were extracted. For whole-mount confocal imaging, the ventral surface of the brain was adhered to a 35 mm culture dish to limit movement, covered in PBS to maintain tissue hydration, and 10 μm step z-stacks collected using a FluoView FV1000 (Olympus) confocal microscope equipped with 5X air objective (Olympus). Image analysis and maximal intensity projections were performed in ImageJ. For mapping glomerular position from whole-mount preparations, OBs from postfixed brains were removed intact and epifluorescence images of the dorsal and, subsequently, medial surfaces of each OB captured at 5x magnification, with each surface positioned roughly perpendicular to the imaging axis. Images were stitched manually in Photoshop before registration of OBs and glomerular position mapping.

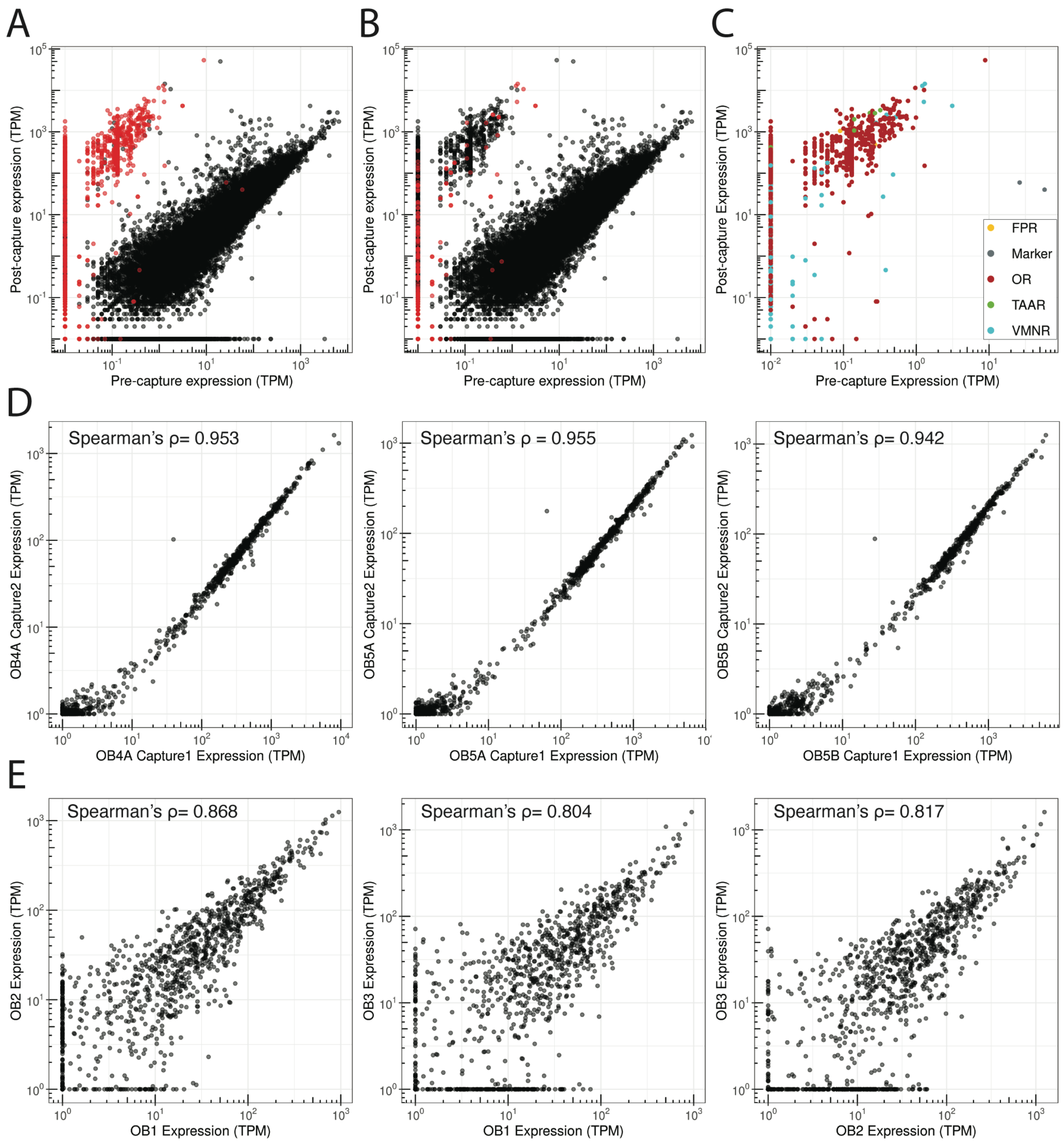


**Fig. S1. Targeted capture sequencing is consistent for enrichment of OR genes.** (**A**) Pre- and post-capture normalized abundances of intact probed genes (red) and intact non-probed genes (black) from a whole OB. (**B**) Pre- and post-capture normalized abundances of intact vomeronasal (Vmnr) genes (red) and intact non-Vmnr genes (black) from a whole OB. (**C**) Pre- and post-capture normalized abundances of target gene families. (**D**) Technical replicates of OR and TAAR gene abundances from independent capture enrichments using two different whole-OB RNA samples (OB4, OB5) combined with different ERCC spike-in mixtures (A, B). (**E**) Biological replicates of OR and TAAR gene abundances from independent capture enrichments of three different whole-OB samples (OB1, OB2, OB3).

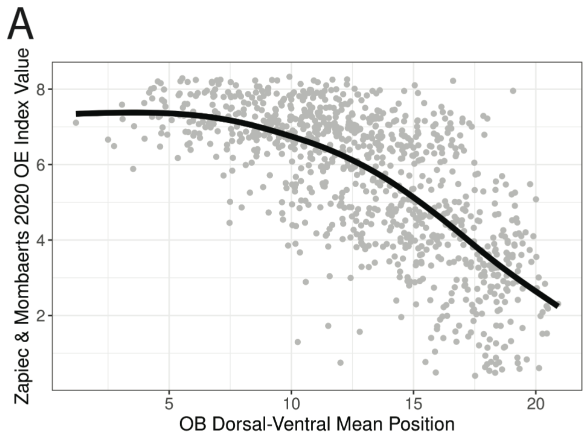


**Fig. S2. Dorsoventral OB spatial sections correlate with known OE positions.** (**A**) Line is loess smoothed regression of OE DV index from Zapiec and Mombaerts, *Cell. Reps.* 2020 across DV mean positions from our targeted spatial data.

A

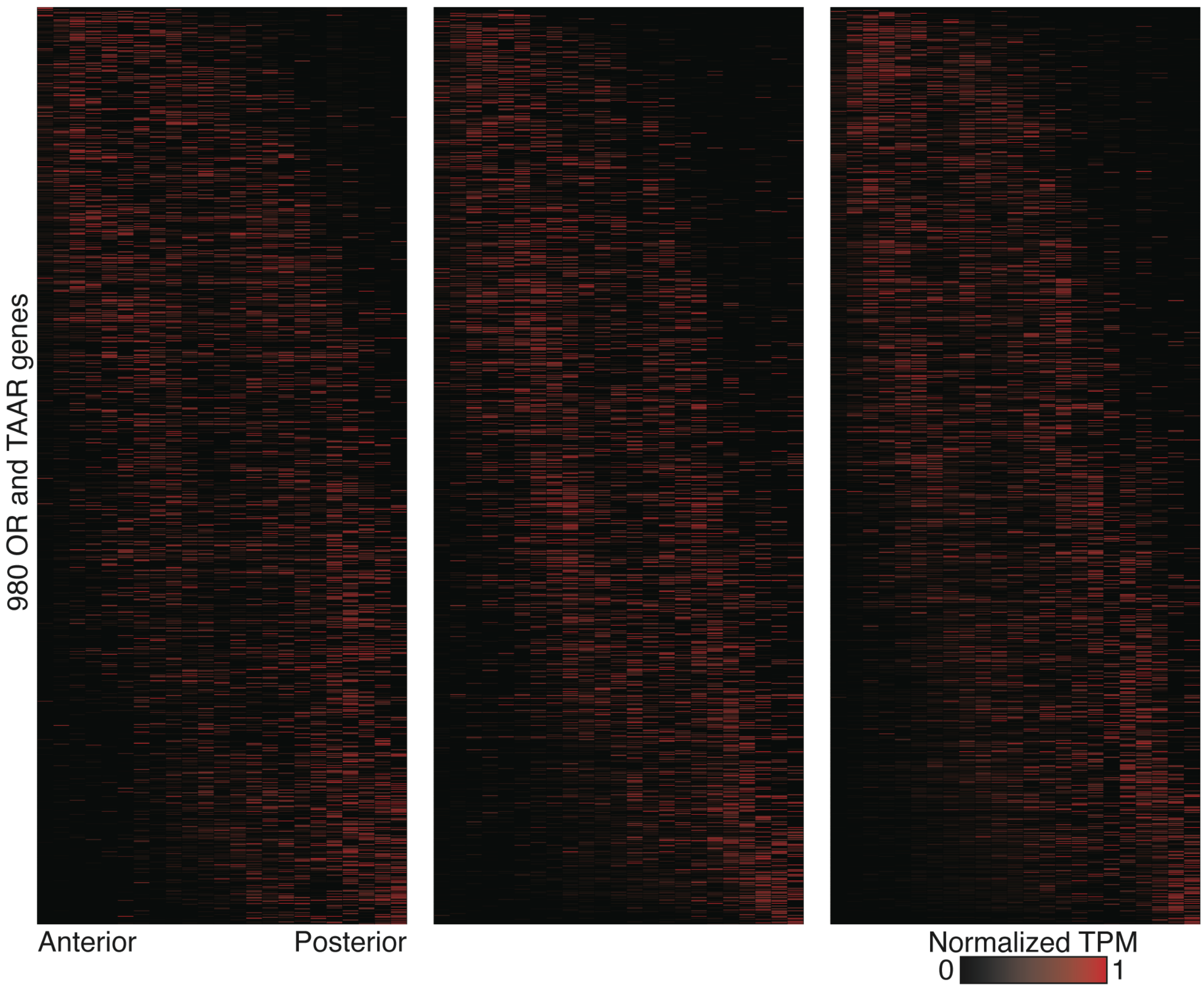


**Fig. S3. ORs display bimodal distributions along the anteroposterior axis.** (**A**) Heatmaps for 980 ORs and TAARs across 23 AP sections sorted by mean position of expression from three additional replicate mice.

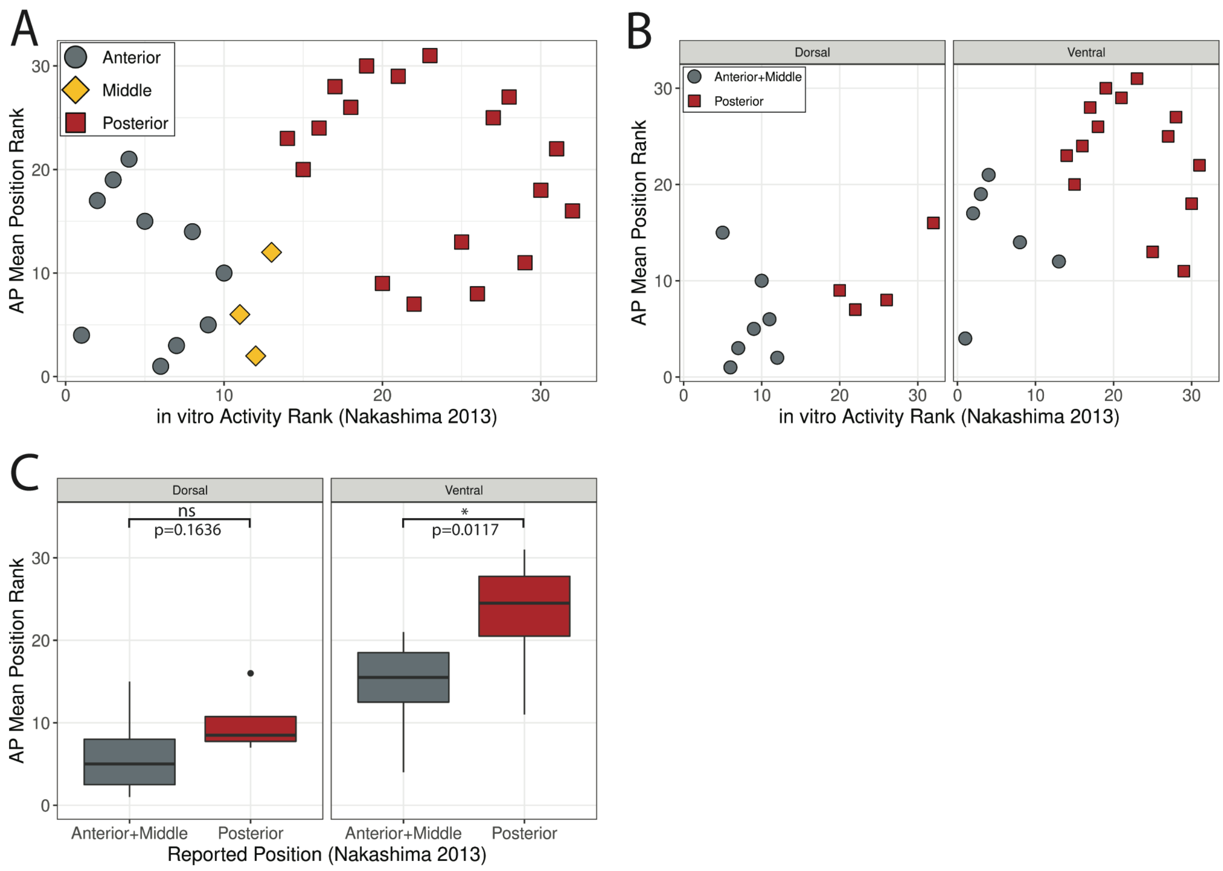


**Fig. S4. Mapped AP positions are consistent with published AP data.** (**A**) Scatter plot of 31 ORs present in our AP dataset examining the relationship between ranked mean AP position and in vitro activity rank from Nakashima et al. *Cell*. 2013. Color indicates the reported OB position from which the OR was cloned from. (**B**) Data from B separated by OE DV zone. (**C**) Data from B, grouped. Statistic is Mann-Whitney U-test.

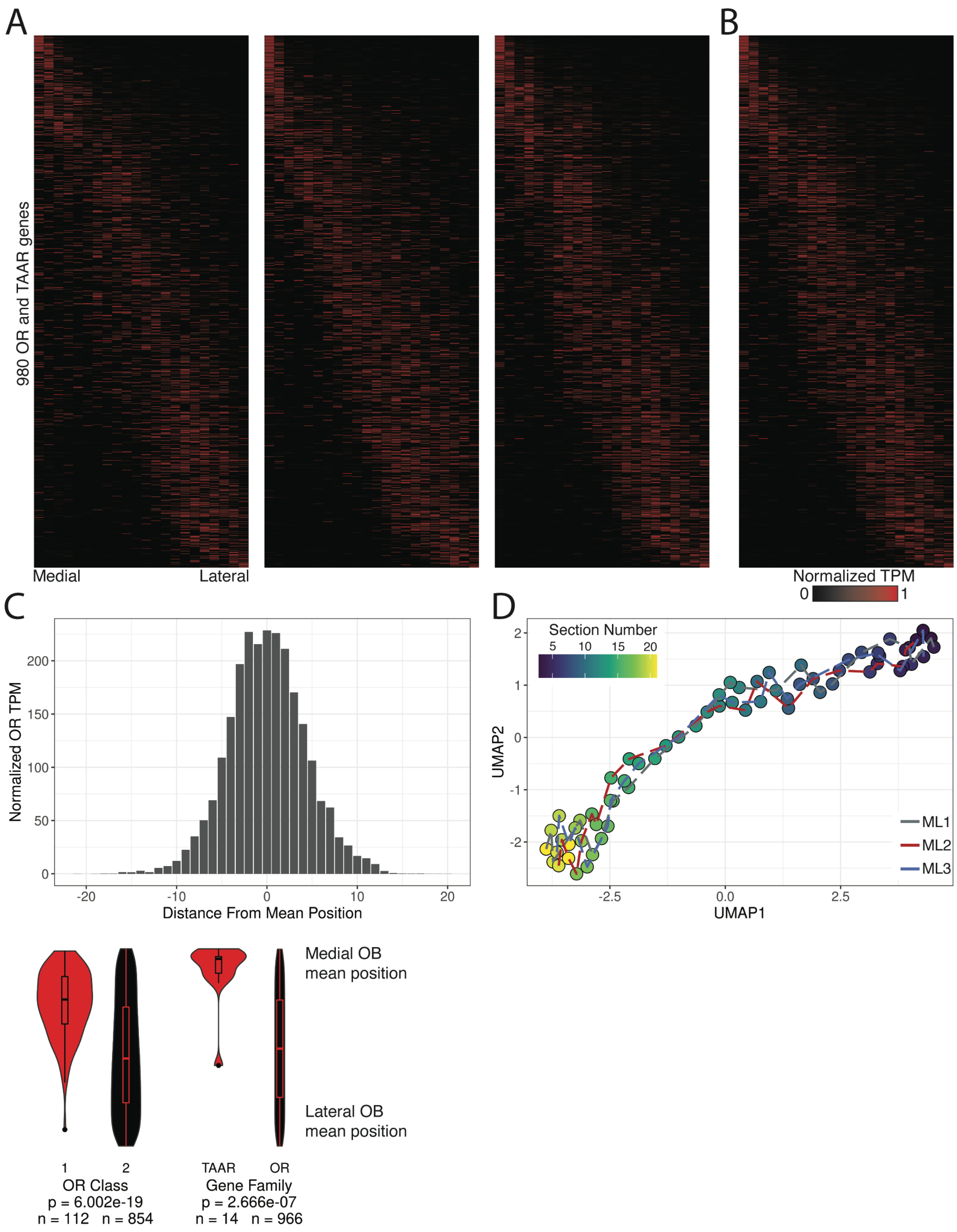


**Fig. S5. Mapping of mediolateral positions.** (**A**) Heatmaps for 980 ORs and TAARs across 22 ML sections sorted by mean position of expression from three replicate mice. (**B**) Merged representation of A. Order of Y-axis genes is consistent across all heatmaps. (**C**) Distribution of normalized TPM (maximum observed value = 1, minimum observed value = 0) for all 980 ORs and TAARs from position of mean expression. (**D**) UMAP projection of 66 ML samples from all three replicates. (**E**) Distribution of ranked ML mean positions for the 980 ORs and TAARs by OR class and gene family. Statistic is Mann-Whitney U-test.

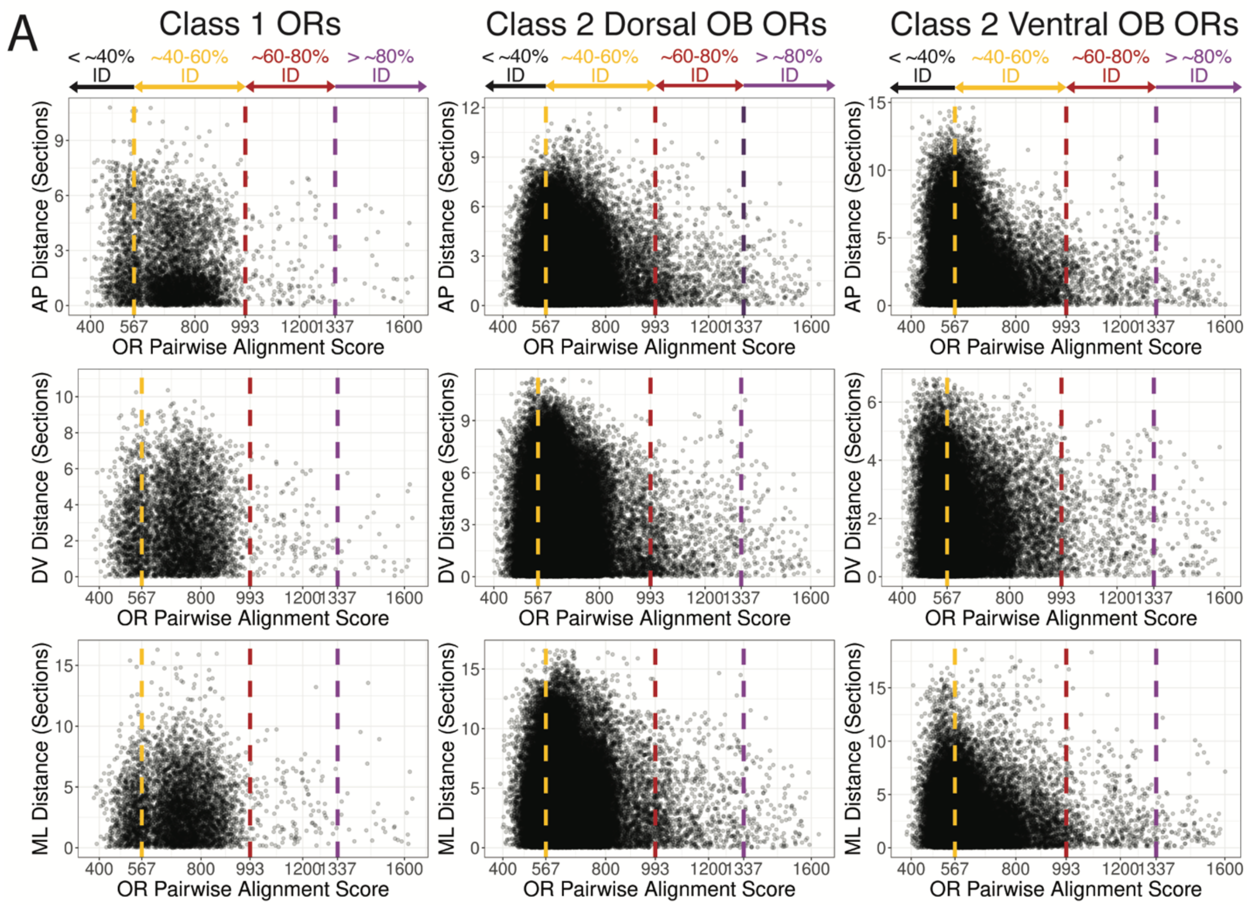


**Fig. S6. Relationship of OR protein sequence and OB position.** (**A**) Scatter plot of mean position distance and OR protein alignment score for all ORs. Score cutoffs are approximations for OR nomenclature values (Family/“567-993” = percent identity ≥ 80%, Subfamily/“993 to 1337” = 80% > percent identity ≥ 60%, Highly similar/“>1337” = percent identity > 80%) for the AP (top), DV (middle), and ML dimensions (bottom).

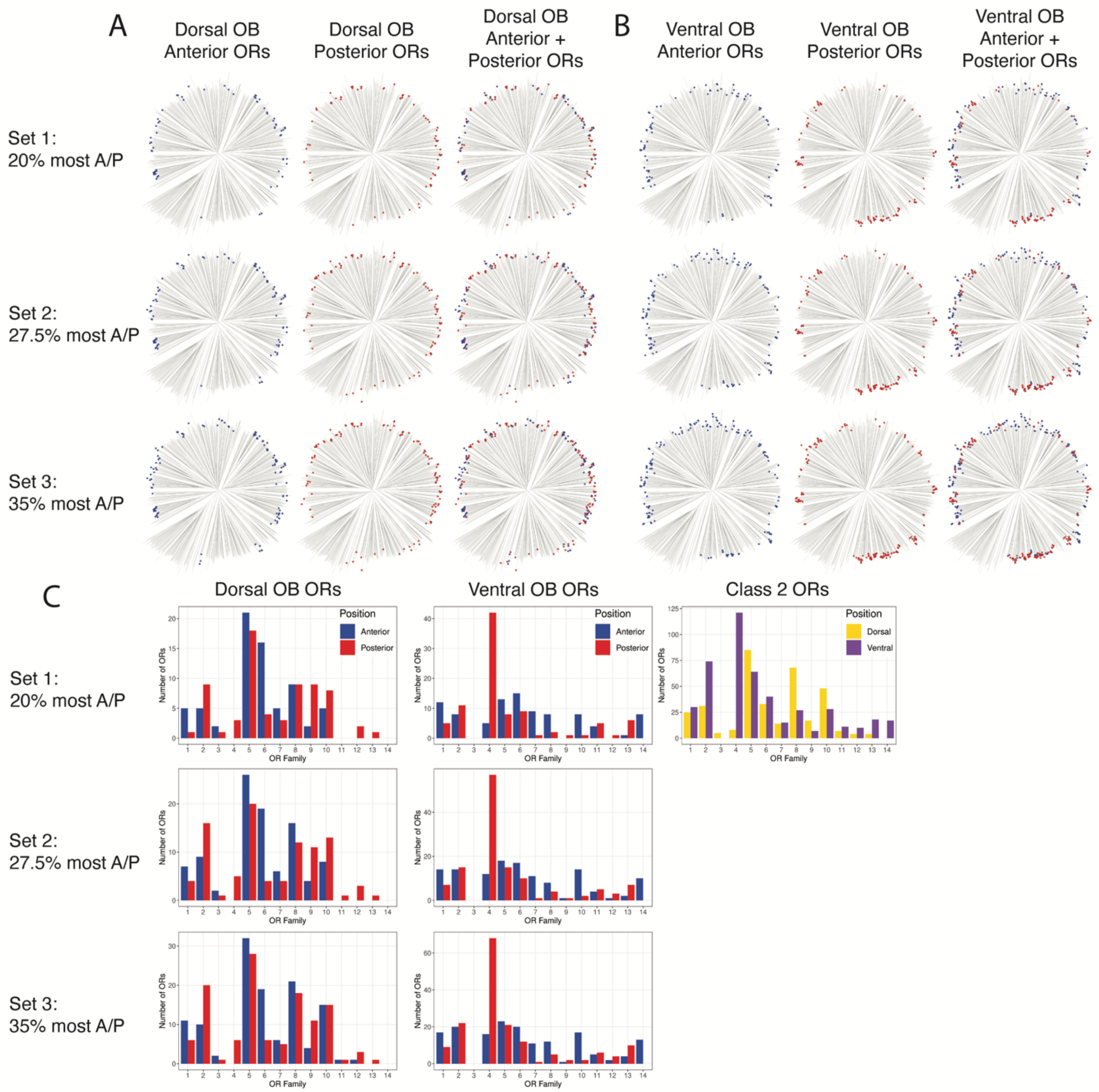


**Fig. S7. Anterior and posterior ORs along the dorsal and ventral OB.** (**A**) Phylogenetic trees for anterior sets of Class II dorsal OB ORs (blue, left), posterior sets of Class II dorsal OB ORs (red, middle), and combined sets of dorsal anterior and posterior Class II OB ORs (right). Sets of different sizes (top row = 20% most anterior/posterior ORs, n = 70; middle row = 27.5% most anterior/posterior ORs, n = 97; bottom row = 35% most anterior/posterior ORs, n = 123) used as conditions for identifying significantly different residues as compared to Class II dorsal OB ORs. (**B**) Phylogenetic trees for anterior sets of Class II ventral OB ORs (blue, left), posterior sets of Class II ventral OB ORs (red, middle), and combined sets of ventral anterior and posterior Class II OB ORs (right). Sets of different sizes (top row = 20% most anterior/posterior ORs, n = 92; middle row = 27.5% most anterior/posterior ORs, n = 127; bottom row = 35% most anterior/posterior ORs, n = 162) used as conditions for identifying significantly different residues as compared to Class II dorsal OB ORs. (**C**) Distributions of dorsal and ventral anterior and posterior ORs by OR family as classified in Olender et al. *BMC Evol. Bio.* 2020 for dorsal OB ORs (left) and ventral OB ORs (middle). Distribution of all Class II dorsal and ventral ORs by OR family is depicted in the top right.

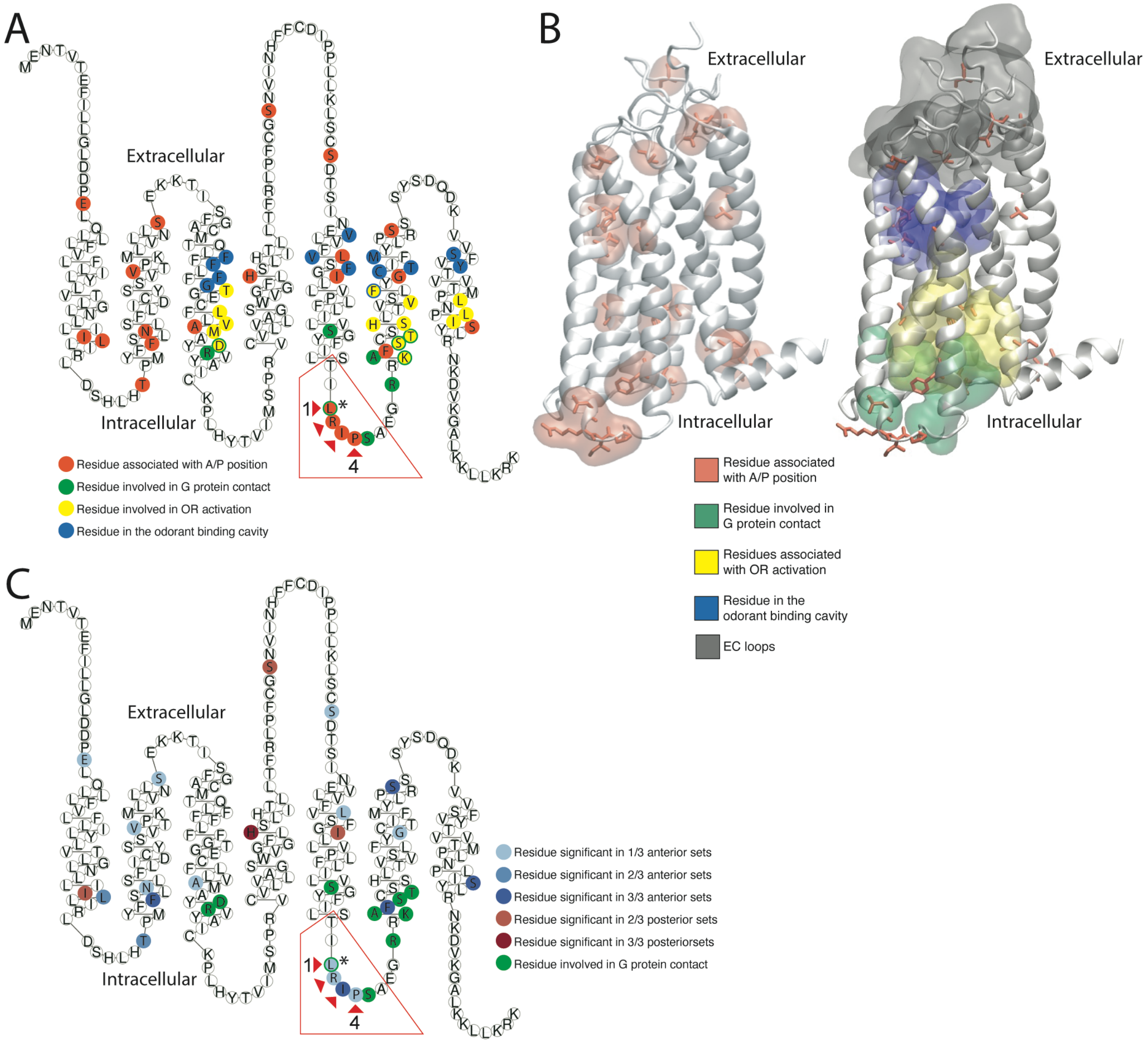


**Fig. S8. Position of anterior and posterior associated residues within ventral Class II ORs.** (**A**) Snakeplot of the Class II OR consensus protein sequence with residues color coded as being associated with glomerular AP position (orange), involved in G protein contact (green), involved in the OR activation mechanism (yellow), and located in the odorant binding cavity (blue). Residues filled with one color and bordered in a different color were associated with both categories. (**B**) Homology model of the mouse Class II consensus OR. Left, residues associated with AP positions (orange) are depicted in licorice with transparent regions indicating residue surface. Right, transparent regions represent the surface of residues located in the extracellular (EC) loops (gray), located in the odorant binding cavity (blue), associated with the activation mechanism (yellow) and associated with G protein binding (green). (**C**) Snakeplot of the Class II OR consensus protein sequence with blue shaded residues were identified as having significant different physicochemical properties for ventral, anterior, Class II ORs compared to all ventral Class II ORs. Red shaded residues were identified as having significantly different physicochemical properties for ventral, posterior, Class II ORs compared to all ventral Class II ORs. Residues highlighted in green indicate mammalian OR residues known to be involved in Class A GPCR activation through contact with the G protein (* indicates the single residue which was identified as being both associated with G protein contact and identified as significantly different for ventral, anterior, Class II ORs).

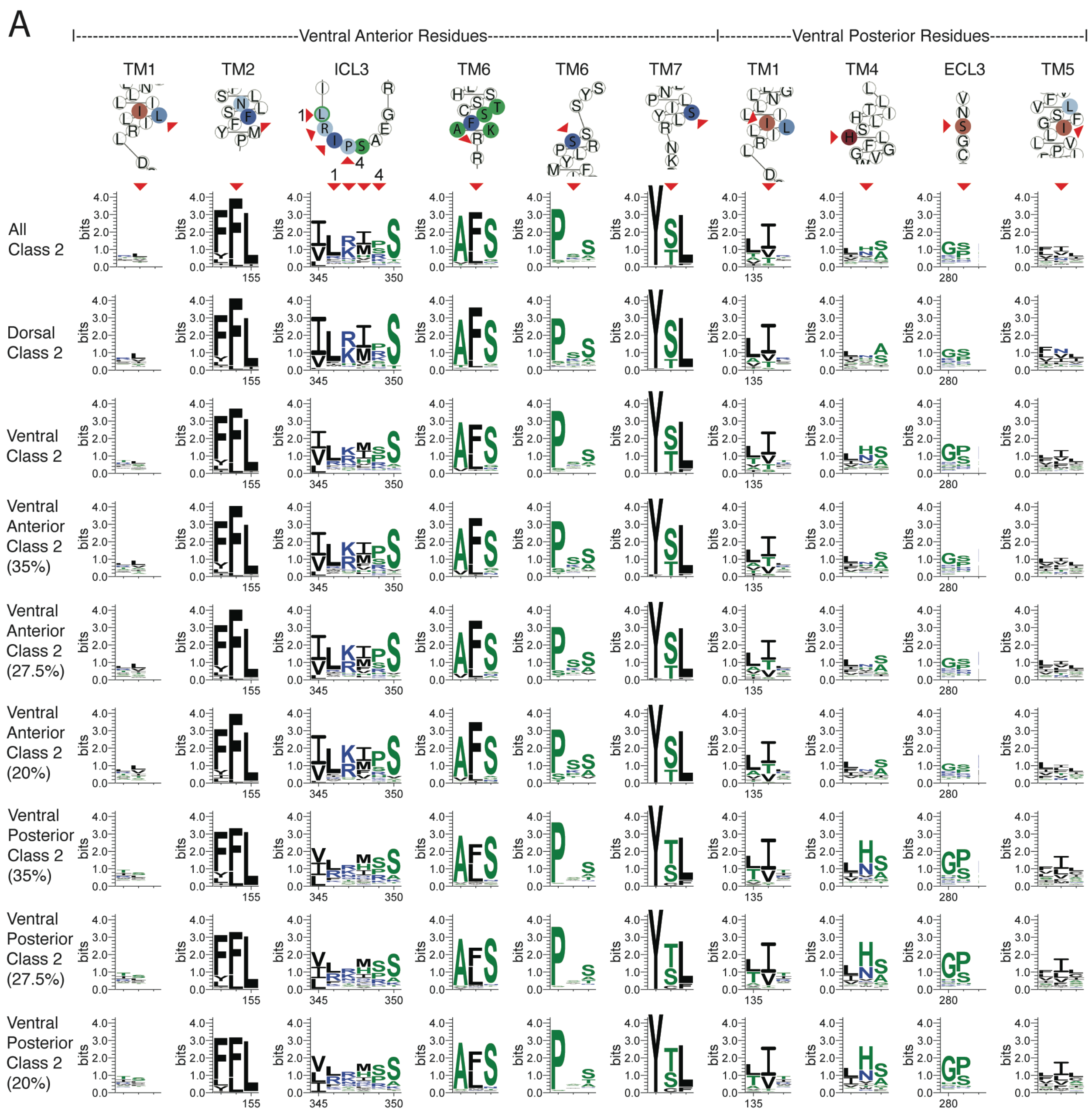


**Fig. S9. Sequence logos for selected ventral Class II anterior and posterior residues.** Protein sequence logos for positions associated with ventral Class II anterior OR residues (left six columns) and ventral Class II posterior OR residues (right four columns) depicting the conservation of specific amino acid residues within different sets of sequences. Red arrows indicate the specific residue within the Class II OR consensus snakeplot (fig. S8, A and C) and the corresponding position in the sequence logo.

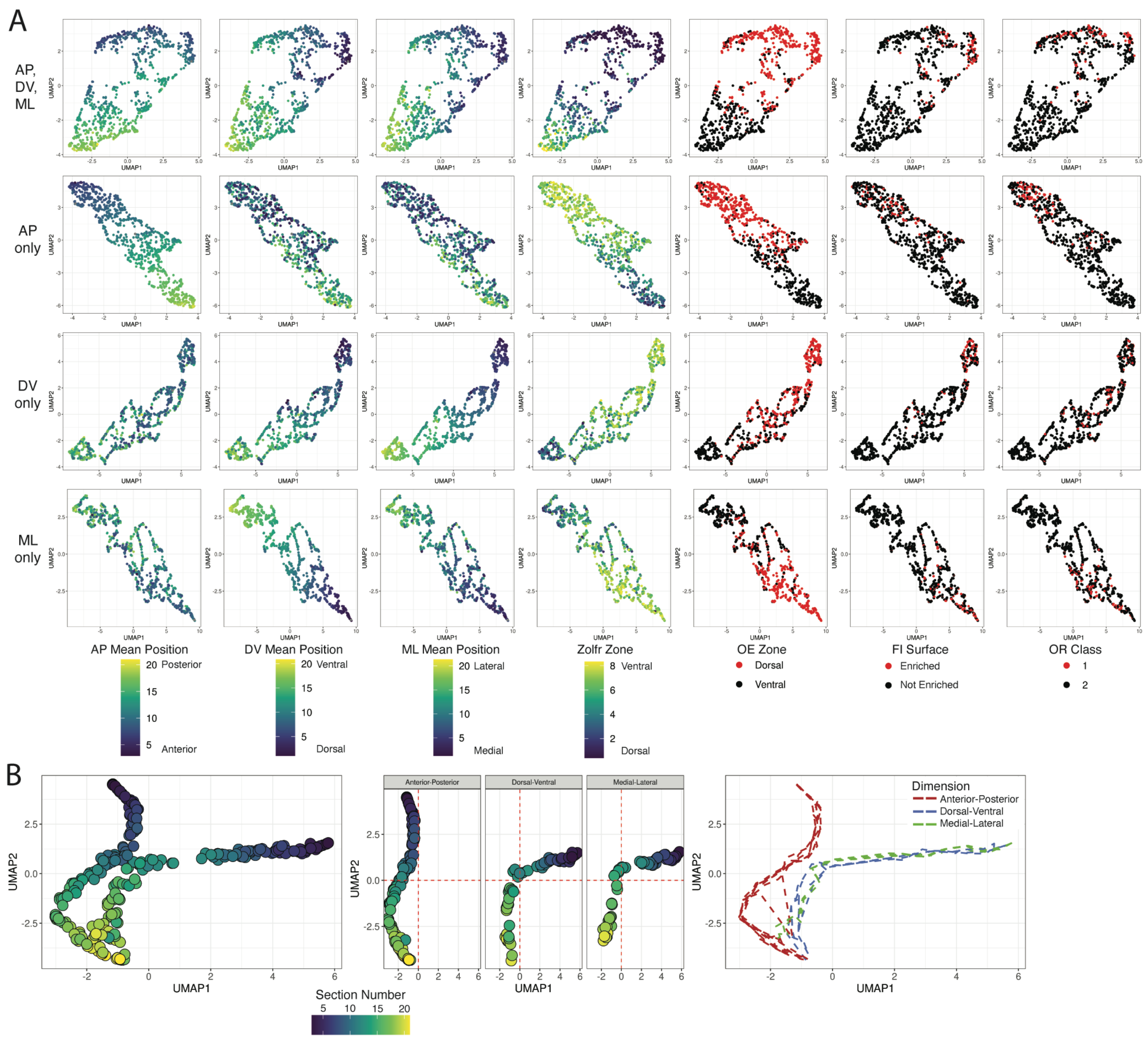


**Fig. S10. Positional features are conserved when only mean position data is used.** (**A**) UMAP projections of OR populations constructed using the mean position from each replicate from all dimensions (top row), AP replicates only (second row), DV replicates only (third row), and ML replicates only (bottom row). AP, DV, and ML mean position reflect calculated OR values for normalized expression weighted means across single dimensions. Zolfr Zone refers to the OE index established by Zapiec and Mombaerts *Cell Reports* 2020. OE Zone refers to the OE index established by Tan and Xie *Chem.*

*Senses* 2018 with values less than 2 labeled as Dorsal and equal or greater than 2 labeled as Ventral. FI Surface refers to the differential expression analysis calculated from dorsal and ventral OB samples in this paper. (**B**) UMAP projection of captured OB spatial samples constructed using transcript abundances for the 980 ORs and TAARs from all dimensions (left), separated by dimension (middle), and for the order of section number by replicate from each dimension (right).

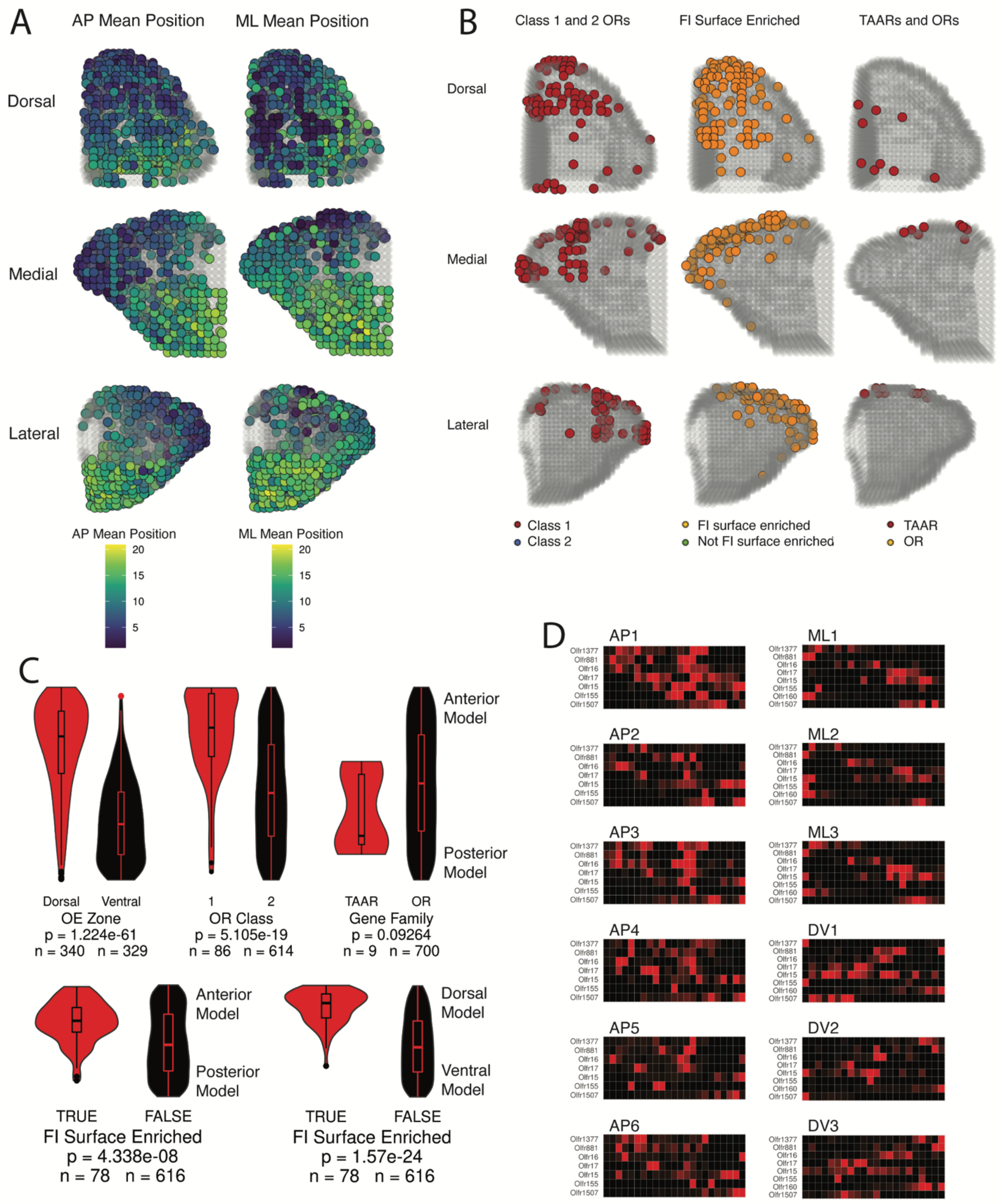


**Fig. S11. A three-dimensional model for OR glomeruli positions from single-dimension targeted sequencing data.** (**A**) Three-dimensional predictions for all 709

ORs and TAARs colored by AP mean position (left) and ML mean position (right). (**B**) Three-dimensional predictions for the 709 ORs and TAARs colored to show the distribution of Class I, functional imaging surface enriched ORs, and TAARs without opposing features in order to better demarcate positions. (**C**) Distribution of ranked mean model position (AP axis for top row and bottom left and DV axis for bottom right) for the best probability voxel in each predicted glomerulus for all 709 ORs and TAARs for OE zone, OR Class, gene family, and enrichment in the functional imaging surface. Statistic is Mann-Whitney U-test. (**D**) Heatmaps for labeled ORs from individual replicates along the AP (left), ML (top right), and DV (bottom right) dimensions.

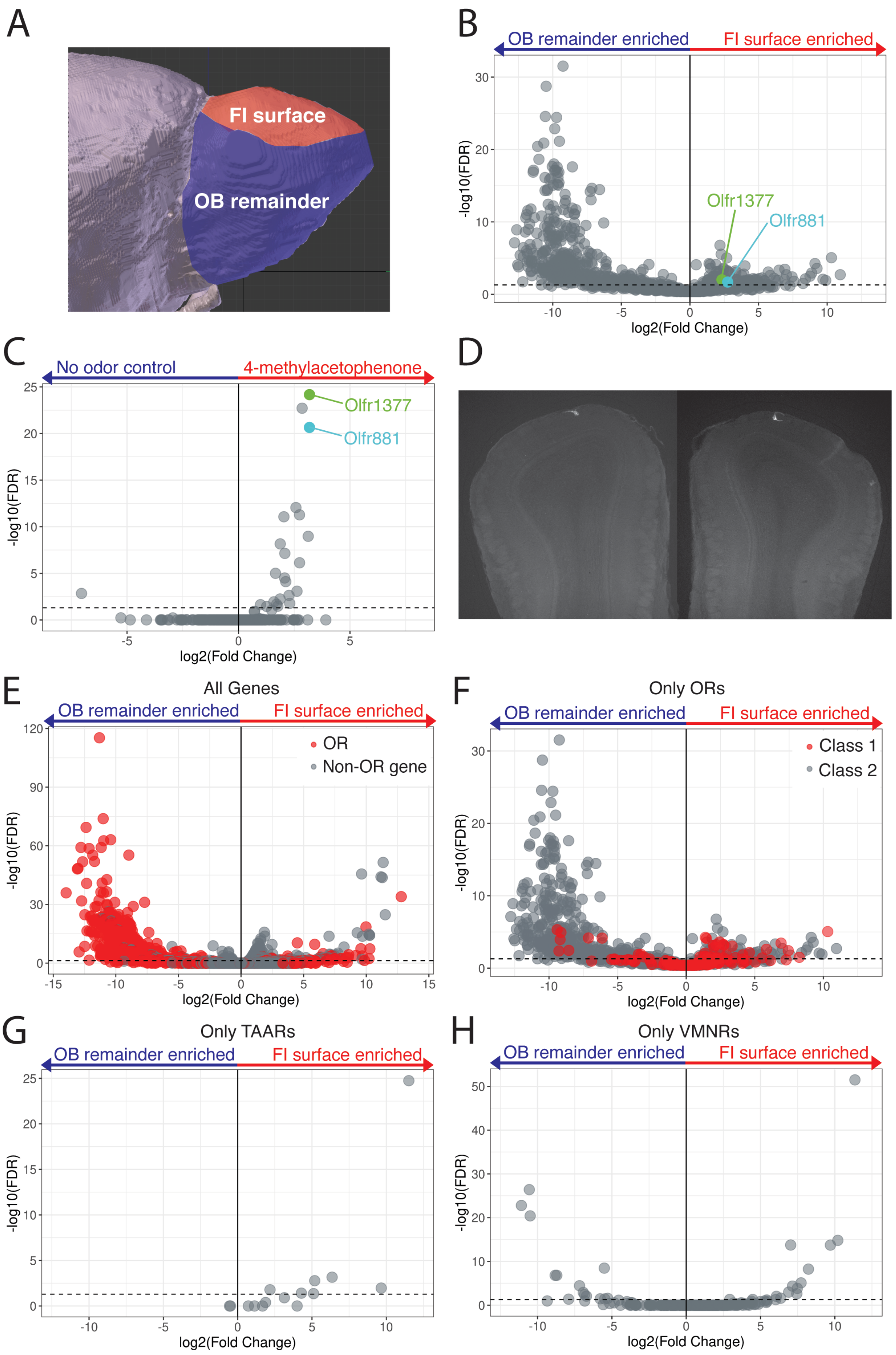
A
FI surface
OB remainder
B
OB remainder enriched
FI surface enriched
Olfr1377
Olfr881
-log10(FDR)
log2(Fold Change)
C
No odor control
4-methylacetophenone
Olfr1377
Olfr881
-log10(FDR)
log2(Fold Change)
D
E
All Genes
OB remainder enriched
FI surface enriched
OR
Non-OR gene
-log10(FDR)
log2(Fold Change)
F
Only ORs
OB remainder enriched
FI surface enriched
Class 1
Class 2
-log10(FDR)
log2(Fold Change)
G
Only TAARs
OB remainder enriched
FI surface enriched
-log10(FDR)
log2(Fold Change)
H
Only VMNRs
OB remainder enriched
FI surface enriched
-log10(FDR)
log2(Fold Change)

372

Fig.

**S12. Differential expression analysis for functional imaging surface samples.** (**A**) OB schematic showing approximation of functional imaging (FI) surface (red) and OB remainder (blue) dissected for differential expression analysis. (**B**) Volcano plot comparing expression of ORs in functional imaging surface and OB remainder samples. The FI surface enriched ORs, Olfr1377 and Olfr881 are labeled in green and cyan, respectively. (**C**) Volcano plot comparing expression of ORs in pS6-IP RNA-Seq stimulation with 1% 4-methylacetophenone. The highly ranked responding ORs, Olfr1377 and Olfr881 are labeled in green and cyan, respectively. (**D**) Coronal sections from an Olfr881-IRES-mKate2 mouse indicating a dorsal-central location of the mKate2 labeled ORs (**E**) Volcano plots for differential expression from functional imaging samples and OB remainder samples for all genes, ORs labeled in red. (**F**) Volcano plot for only ORs, FDR readjusted, Class I ORs labeled in red. (**G**) Volcano plot for only TAARs, FDR readjusted. (**H**) Volcano plot for only vomeronasal receptors (VMNRs), FDR readjusted.

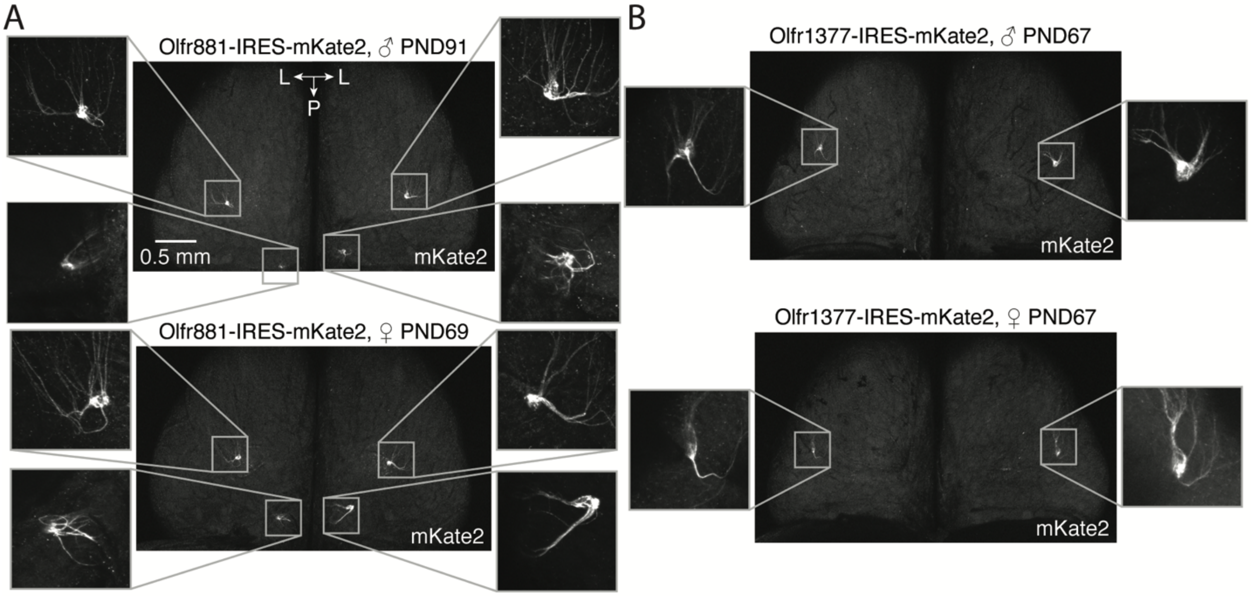


**Fig. S13. Whole-mount confocal microscopy and glomerular phenotypes of Olfr1377-IRES-mKate2 and Olfr881-IRES-mKate2 OBs.** (**A**) Whole-mount maximum intensity projections of the dorsal surface of Olfr881-IRES-mKate2 OBs from male and female mice with inlays showing the mKate2-positive glomeruli. (**B**) Whole-mount maximum intensity projections of the dorsal surface of Olfr1377-IRES-mKate2 OBs from male and female mice with inlays showing the mKate2-positive glomeruli.

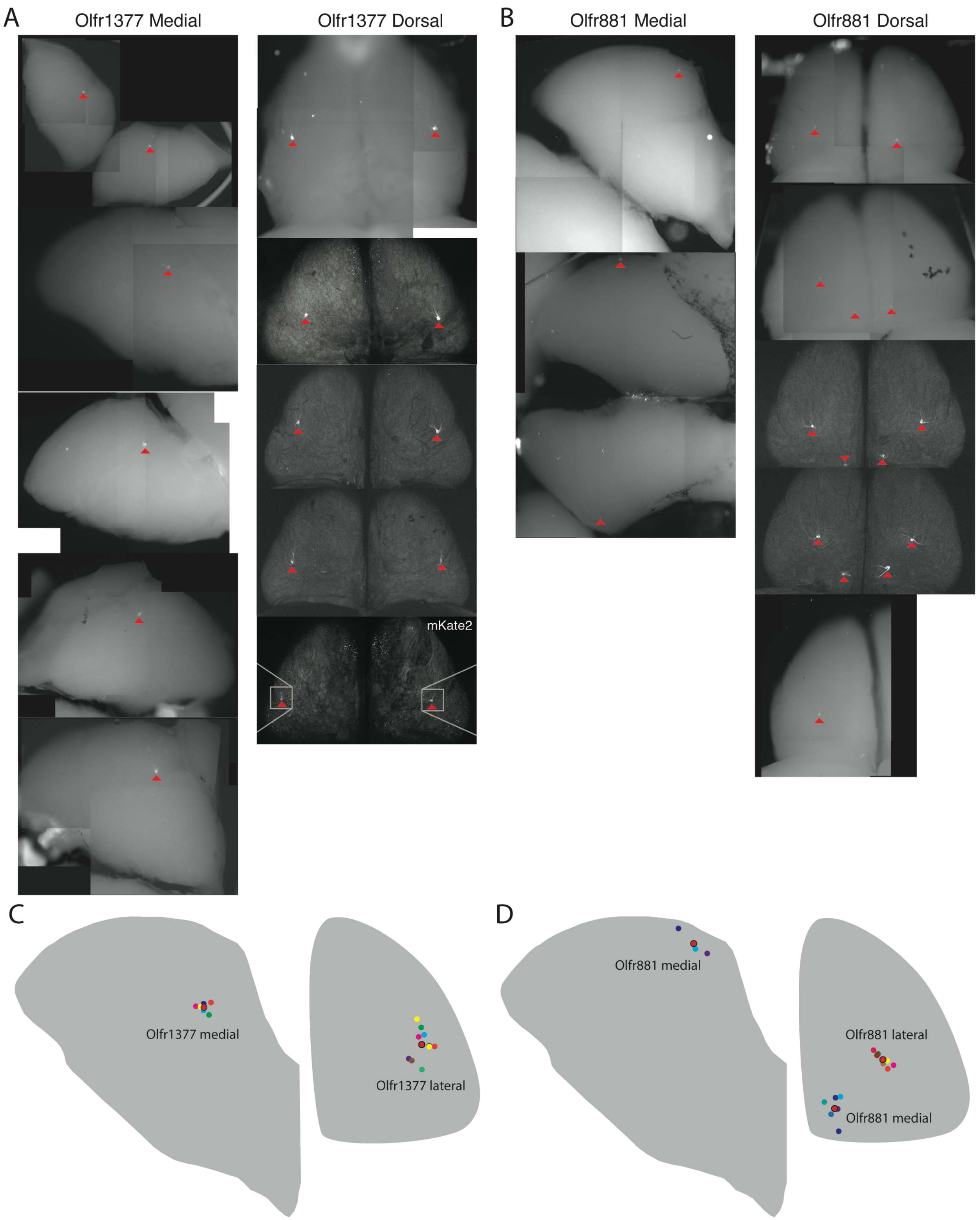


**Fig. S14. Position of Olfr1377 and Olfr881 glomeruli.** (**A**) Whole-mount stitched epifluorescence images (together with whole-mount confocal maximal intensity projections from fig. S13) reconstructing the medial (left) and dorsal (right) faces of Olfr1377-IRES-mKate2 mice OBs. Red arrows indicate glomerulus. (**B**) Same as (A) for Olfr881-IRES-mKate2 mice. (**C**) Positions of Olfr1377 fluorescent glomeruli as recorded from the whole-mount OB images. Red bordered circles indicate the mean position of the individual replicates. (**D**) Same as (C) for Olfr881.

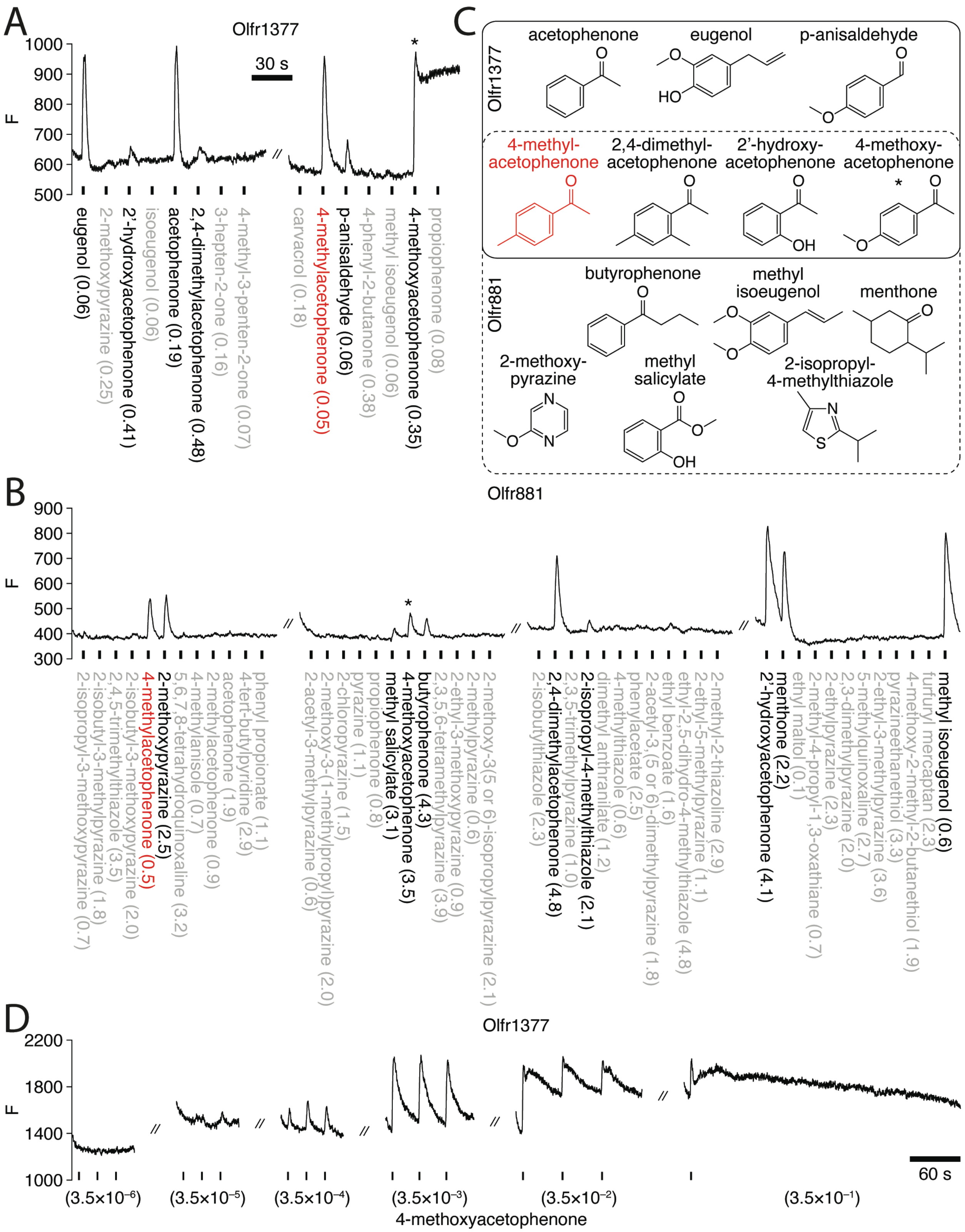


**Fig. S15. In vivo two-photon GCaMP6s response kinetics and concentration-**

**dependence of Olfr1377 and Olfr881 glomeruli.** (**A**) Raw two-photon GCaMP6s fluorescence of a mKate2-labeled Olfr1377 glomerulus during sequential presentation of odorants in pseudorandom order to a compound heterozygous Olfr1377-IRES-mKate2; OMP-IRES-tTA; tetO-GCaMP6s mouse. * marks long-term activation by 4-methoxyacetophenone. Red text highlights response to 4-methylacetophenone. Odorants presented at an estimated concentration on the order of $10^{-1}$ nM. (**B**) Similar to (A) for a mKate2-labeled Olfr881 glomerulus imaged in a compound heterozygous Olfr881-IRES-mKate2; OMP-IRES-tTA; tetO-GCaMP6s mouse. Note the lack of long-term activation by 4-methoxyacetophenone. Odorants presented at an estimated concentration on the order of $10^{0}$ nM. (**C**) Chemical structures of a subset of ligands identified by functional imaging for Olfr1377 (solid box) and Olfr881 (dashed box), including multiple overlapping cyclic ketone ligands. (**D**) Raw two-photon GCaMP6s fluorescence of a mKate2-labeled Olfr1377 glomerulus during sequential presentation of increasing concentrations of 4-methoxyacetophenone, revealing long-term activation with $3.5 \times 10^{-2}$ nM and higher concentrations.

### SUPPLEMENTARY TABLES

**Table S1**: Mice, sex, age, and number of samples

**Table S2**: Sequencing samples and read depth

**Table S3**: Target genes, probe coverage, fold enrichment

**Table S4**: Mean positions for individual and merged replicates along the AP, DV, and ML dimensions for each OR

**Table S5**: Summary values and statistics for percent identity score-based groups of Class I, Class II Dorsal OB ORs, and Class II Ventral OB ORs

**Table S6**: Residue identities of Class II OR consensus sequence and significance for all tested sets of anterior/posterior ORs

**Table S7**: Three-dimensional model inputs, weights, posterior probabilities, predicted locations

**Table S8**: Differential expression analysis for functional imaging surface and pS6-IP 4-methylacetophenone studies

**Table S9**: Heterologous cell assay results
